# Dancing with the ions: symport and antiport mechanisms of human glutamate transporters

**DOI:** 10.1101/2022.08.12.503774

**Authors:** Biao Qiu, Olga Boudker

**Affiliations:** Department of Physiology & Biophysics, Weill Cornell Medicine, 1300 York Ave, New York, NY 10021, USA; Howard Hughes Medical Institute, Weill Cornell Medicine, 1300 York Ave, New York, NY 10021, USA

**Keywords:** Glutamate transporter, ions coupling, symport and antiport

## Abstract

Excitatory amino acid transporters (EAATs) pump glutamate into glial cells and neurons. EAATs achieve million-fold transmitter gradients by symporting it with three sodium ions and a proton and counter-transporting a potassium ion via an elevator mechanism. Despite the availability of structures, the symport and antiport mechanisms remain unclear. We report high-resolution Cryo-EM structures of human EAAT3 bound to the neurotransmitter glutamate with symported ions, potassium ions, sodium ions alone, or in the absence of ligands. We show that an evolutionarily conserved occluded translocation intermediate has a dramatically higher affinity for the neurotransmitter and the counter-transported potassium ion than outward- or inward-facing transporters and plays a crucial role in ion coupling. We propose a comprehensive ion coupling mechanism involving a choreographed interplay between bound solutes, conformations of conserved amino acid motifs, and movements of the gating hairpin and the substrate-binding domain.

## Introduction

Human excitatory amino acid transporters (EAATs) concentrate neurotransmitter glutamate and aspartate in cells (Arriza et al., 1997; Fairman et al., 1995; Kanai and Hediger, 1992; Pines et al., 1992; Storck et al., 1992). By coupling uptake of one substrate molecule to movements of five ions - symport of three sodium (Na^+^) ions and a proton and counter-transport of a potassium (K^+^) ion - EAATs ensure that the cytoplasmic glutamate concentration is ∼10 mM, required for neuronal functions, while the extracellular concentrations are as low as a few nM permitting rounds of synaptic signaling (Herman and Jahr, 2007; Levy et al., 1998; Owe et al., 2006; Zerangue and Kavanaugh, 1996a). Dysfunction of EAATs leads to dysregulation of glutamate concentrations associated with central nervous system pathologies such as epilepsy, neurodegeneration, and dicarboxylic aminoaciduria (Bailey et al., 2011; Berman et al., 2011; Duerson et al., 2009; Jen et al., 2005; Masliah et al., 1996; Sepkuty et al., 2002; Tanaka et al., 1997).

EAATs are homotrimers composed of promoters working independently (Grewer et al., 2005; Koch et al., 2007; Leary et al., 2007). TMs 1, 2, 4, and 5 of each protomer comprise the scaffold domain mediating trimerization; TMs 3, 6, 7, and 8, and helical hairpins (HPs) 1 and 2 form the substrate-binding transport domain (Figure S1A). EAATs transport glutamate and aspartate with similar apparent affinities (Arriza *et al*., 1997; Arriza et al., 1994; Fairman *et al*., 1995; Storck *et al*., 1992) by an elevator mechanism, whereby the substrate-bound transport domain rotates and translocates over 15 Å across the membrane from an outward- to an inward-facing state (OFS and IFS, Figure S1B and Movie S1) (Canul-Tec et al., 2017; Canul-Tec et al., 2022; Qiu et al., 2021; Zhang et al., 2022). EAATs also mediate an uncoupled anion flux (Billups et al., 1996; Eliasof and Jahr, 1996; Fairman *et al*., 1995; Vandenberg et al., 1995; Wadiche et al., 1995), where the conducting state is likely a transient intermediate of the elevator transition (Chen et al., 2021; Cheng et al., 2017; Machtens et al., 2015; Pant et al., 2022).

The EAATs’ transport cycle begins with the transport domain in OFS, where it binds sodium ions, a proton, and the transmitter. The transport domain moves into IFS, releases the solutes, binds a potassium ion, and translocates it back into OFS for release into the extracellular space. Earlier studies suggest that cooperative binding of the transmitter, sodium ions, and protons underlies coupled transport (Arkhipova et al., 2020; Boudker et al., 2007; Guskov et al., 2016; Oh and Boudker, 2018; Qiu *et al*., 2021; Verdon et al., 2014; Wang and Boudker, 2020b). Two conserved “dancing” motifs rearrange in response to sodium binding, increasing substrate affinity (Figure S1C): NMD_366-368_ in the unwound region of TM7 and YE_373-374_ in TM7b and DxxxDxxR_440-447_ in TM8 (NMD and YE/DDR motifs for short). In the absence of Na^+^ ions, the motifs are in the “apo” configurations: the M367 sidechain points out of the substrate-binding site, and R447, salt-bridged to deprotonated proton carrier E347, occupies the site (Qiu *et al*., 2021). Sodium binding to Na1 and Na3 sites induces the “bound” configuration: M367 swings into the binding pocket, poised to coordinate the third Na^+^ ion in the Na2 site (Zhou et al., 2022). R447 disengages from E347 and swings out of the pocket, forming cation-π interactions with Y373 and ready to coordinate the substrate. E347 binds a proton. The consequent binding of the transmitter and Na2, or the binding of K^+^ ions to the apo transporter, is thought to close HP2, allowing elevator movements (Arkhipova *et al*., 2020; Boudker *et al*., 2007; Canul-Tec *et al*., 2022; Guskov *et al*., 2016; Qiu *et al*., 2021; Verdon *et al*., 2014; Wang and Boudker, 2020b). The energetic penalty for rearranging the motifs into the bound configuration underlies the cooperative binding of the transmitter and ions.

Here, we describe new Cryo-EM structures of human neuronal and epithelial EAAT3 (hEAAT3), which, together with our earlier structures (Qiu *et al*., 2021), led us to overhaul the ion-coupling hypotheses of glutamate transporters.

We propose that an evolutionarily conserved transient occluded intermediate outward-facing state (iOFS) ensures coupled transport by discriminating between translocation competent and incompetent transport domains. In hEAAT3, iOFS might be the only state that binds the transmitter and potassium tightly. In the translocation-competent substrate/3Na^+^/proton- and K^+^-bound states, the “zipped” gating HP2 completely occludes the substrate and ion binding cavities. In contrast, the distorted hairpin is “unzipped” in the translocation-incompetent partially bound and apo states, allowing water access to the substrate- and ion-binding sites. Proton binding to E374 favors the transition from iOFS to OFS, the release of potassium ion, the opening of HP2, and the isomerization of the dancing motifs into the bound-like configuration, priming the transporter for sodium and transmitter binding. Potassium binds to the transporter with the apo dancing motifs in the place of the substrate amino group, sequestering a coordinating residue from the Na2 site. Thus, the structures explain the strictly competitive binding of the symported glutamate/3Na^+^/H^+^ and the counter-transported potassium ions. The structures also reveal that the evolutionary molecular innovation, leading to the expansion of ion coupling in eukaryotes to include potassium ions and protons, relies on the glutamine to glutamate replacement in YE/DDR motif. In archaeal transporters, glutamine stabilizes the zipped conformation of HP2, while in eukaryotes, either protonation of E374 or potassium binding is required.

Our structural analyses provide a complete description of the transport cycles of hEAAT3 and suggest the energetic relationships between the states, which ensure coupling of the neurotransmitter uptake the ions fluxes down their electrochemical gradients.

## Results

### Intermediate occluded states of hEAAT3 feature closed HP2 gates

In earlier studies, we could not image potassium- or glutamate-bound hEAAT3 because the transporter did not bind the solutes well in its preferred IFS (Qiu *et al*., 2021). Thus, we aimed to constrain hEAAT3 in OFS by cysteine crosslinking, expecting the state to show higher affinities. First, we prepared hEAAT3 with mutated glycosylation sites (hEAAT3g) (Qiu *et al*., 2021) and minimal cysteines to reduce spurious crosslinking. Sequence alignments showed that four of six EAAT3 cysteines are not conserved; the fifth is tryptophan in all other EAAT homologues (Qiu *et al*., 2021). Mutating these cysteines in hEAAT3g yielded MinCys EAAT3 with a size exclusion chromatography (SEC) profile similar to hEAAT3g (Figure S2A) and robust substrate uptake (Figure S2B). However, even conservative mutations of the sixth highly conserved C343 in HP1 yielded denatured inactive proteins (Figure S2A, B). We then introduced K269C and W441C mutations in the scaffold and transport domains of MinCys EAAT3, respectively, because these residues crosslink in OFS in cell-based assays (Shabaneh et al., 2014; Wang et al., 2022). Notably, C343 is distant and unlikely to crosslink.

We crosslinked MinCys EAAT3 K269C/W441C using Hg^2+^ (EAAT3-X) and imaged apo transporter in 150 mM N-methyl-D-glucamine (NMDG) chloride, sodium-bound transporter in 300 mM NaCl, neurotransmitter-bound transporter in 200 mM NaCl and 20 mM glutamate, and potassium-bound transporter in 300 mM KCl by Cryo-EM. Together, these structures should describe the sequential binding of coupled solutes and visualize the potassium counter-transport. We first refined all EM maps applying C3 symmetry and then performed symmetry expansion followed by local 3D classification. In each dataset, we found transporter protomers in OFS and a conformation resembling iOFS of archaeal homologues Glt_Ph_ and Glt_Tk_ (Arkhipova *et al*., 2020; Verdon and Boudker, 2012; Wang and Boudker, 2020b). Thus, we obtained the EM maps of the dominant conformations in C3 with 2.44-2.8 Å overall resolution and single protomers in the minor conformations with 2.94-3.4 Å resolution (Table S1-3, Figure S3, 4). EAAT3-X was predominantly in OFS in NaCl buffer and iOFS in NMDG chloride, NaCl/glutamate, and KCl buffers (Figure 1). Notably, we observed no K^+^ ions bound in OFS determined in KCl buffer, and assigned the state as OFS-Apo_K_ (described in detail below).

**Figure 1.**
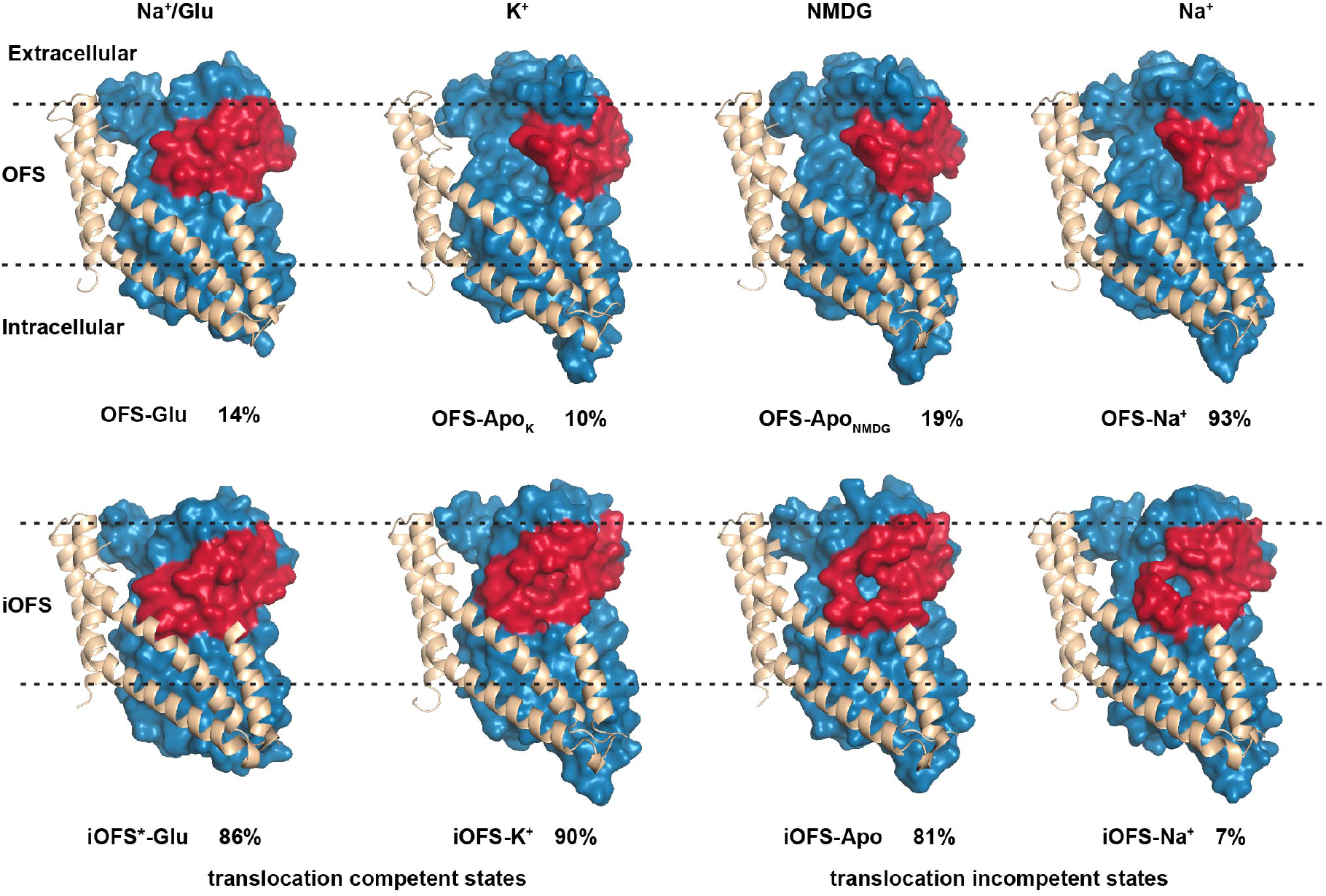
Outward-facing and occluded intermediate states of EAAT3-X. OFS (top) and iOFS (bottom) protomers of EAAT3-X in, from left to right, 200 mM NaCl and 20 mM glutamate (Na^+^/Glu), 300 mM KCl (K^+^), 150 mM NMDG chloride (NMDG), and 300 mM NaCl (Na^+^). Scaffold TMs 1, 2, 5, and part of TM4 are shown as cartoons and colored wheat. The transport domains are shown in surface representation and colored dark blue, with HP2 red. The assigned conformational states and the fractions of protomers found in these states in 3D classifications following symmetry expansion are below the structures. HP2 in iOFS^*^-Glu, OFS-Glu, and iOFS-K^+^ fully occludes the ligand cavity; the cavities of iOFS-Apo and iOFS-Na^+^ remain solvent-accessible; HP2 of OFS-Apo in NMDG chloride and KCl and OFS-Na^+^ are wide-open.

To ensure symport and antiport of ions, the transport domain must undergo elevator transitions either when bound to glutamate/3Na^+^/proton or K^+^ ions and not when apo or bound to sodium only (Figure 1). Thus, we expected translocation-incompetent species to feature open HP2 gates seen in archaeal transporters (Arkhipova *et al*., 2020; Boudker *et al*., 2007; Verdon *et al*., 2014; Wang and Boudker, 2020a). Indeed, all OFS structures showed wide-open HP2 gates, except when bound to glutamate, which closed the gate. However, surprisingly, all iOFS structures showed closed hairpins with tips engaging the scaffold domain. Compact hairpins completely occluded the binding pockets in glutamate/3Na^+^/proton- or K^+^-bound transporters but showed solvent-accessible openings in apo and sodium-bound species (Figure 1).

### Glutamate mediates local and global conformational shifts

The transport domains of the predominant glutamate-bound iOFS shift more inward by ∼3 Å and rotate by an additional 10 °, compared to the iOFS of the archaeal homologues; we named this conformation iOFS^*^-Glu (Figure 2A). The distances between the Cα atoms of C269 and C441 are 9.1 and 7.7 Å in OFS-Glu and iOFS^*^-Glu, respectively. There is excess density connecting the sulfur atoms attributed to Hg^2+^ ions in both structures (Figure S5A, B), consistent with crosslinked cysteines. Thus, crosslinking allows transport domain movements between OFS and iOFS^*^. The domains superimpose with an overall RMSD of 0.74 Å, but the density of the HP2 tip is less well resolved in OFS-Glu than iOFS^*^-Glu, suggesting increased dynamics (Figure S6).

**Figure 2.**
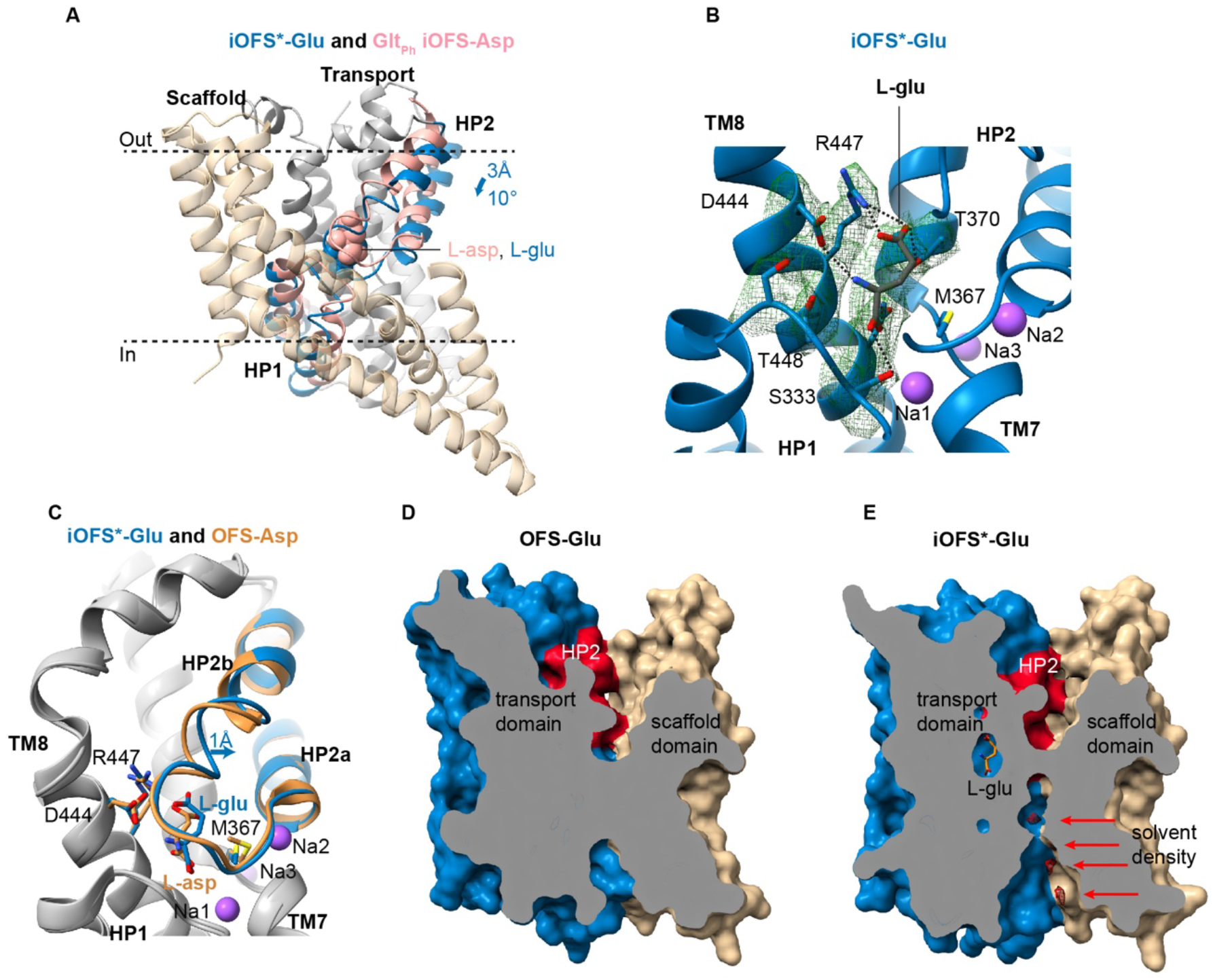
Glutamate binding to EAAT3. (**A**) Superposition of EAAT3-X iOFS^*^-Glu and aspartate-bound Glt_Ph_ iOFS-Asp (PDB ID: 6uwl) aligned on the oligomerization regions of the scaffold domains (tan). The transport domains are colored gray, and HP1 and HP2 are blue in EAAT3-X and pink in Glt_Ph_. Bound L-asp and L-glu are shown as pink and blue spheres. (**B**) Glutamate coordination by EAAT3-X. Bound substrate and coordinating residues are shown as sticks and colored by atom type. Green mesh is the EM density contoured at 5 α. Dashed lines represent hydrogen bonds between L-glu and the coordinating residues. (**C**) Transport domains of aspartate-bound hEAAT3g OFS (OFS-Asp, PDB ID 6×2z) and iOFS^*^-Glu superimposed on their cytoplasmic halves (residues 314-372 and 442-465). Transport domains are colored gray with HP2, substrate, and coordinating residues colored orange and blue, respectively. Surface representation of OFS-Glu (**D**) and iOFS^*^-Glu (**E**), sliced through the substrate-binding site and the domain interface. Arrows emphasize the observed solvent EM densities, shown as red mesh and contoured at 5 σ.

Well-resolved glutamate is coordinated by the highly conserved D444 and R447 in TM8, interacting with the amino group and the sidechain carboxylate, respectively, and S333 in HP1 and N451 in TM8, interacting with the main-chain carboxylate (Figure 2B). Main-chain carbonyl oxygens and amide nitrogens in HP1 and HP2 also contribute to coordination. Overall, glutamate binds similarly to aspartate, except the binding pocket enlarges slightly due to HP2b outward movement and local sidechain shifts (Figure 2C and Movie S2). Interestingly, earlier hEAAT3g structure showed an unusual rotameric state of bound aspartate. In contrast, bound glutamate features the same isomer as aspartate in the archaeal homologs or tsEAAT1 (Boudker *et al*., 2007; Canul-Tec *et al*., 2017; Guskov *et al*., 2016). Thus, glutamate transporters can adjust the substrate-binding mode through local tuning of sidechain conformations and global HP2 movements.

The interface between the transport and scaffold domains is well-packed in OFS-Glu (Figure 2D). In contrast, in iOFS^*^-Glu, a deep crevice between the domains extends from the cytoplasmic side (Figure 2E). There are two constrictions – a tighter one near the extracellular surface and another one closer to the cytoplasm. The cytoplasmic portion of the crevice is similar to a recently proposed chloride-conducting pathway (Chen *et al*., 2021) but distinct from a competing model (Machtens *et al*., 2015). S74 in TM2, previously implicated in the selectivity of the anion channel in EAATs (Cater et al., 2014; Ryan and Mindell, 2007; Ryan et al., 2004), contributes to the cytoplasmic constriction and hydrogen-bonds to a solvent molecule in the crevice. The extracellular constriction, comprised of transport domain residues in TM7 and the HP2 tip (T364, A408, A409, and G410) and scaffold residues in TMs 2 and 4 (I67, I71, V212), has a pore radius of about 0.91 Å suggesting it is closed (Figure S5C, D). We found several excess density peaks in the crevice, including between the constrictions, and modeled them as water molecules because the resolution is insufficient to distinguish water and ions (Figure 2E). We suggest that iOFS^*^ represents a state preceding channel opening, consistent with the more inward position of the transport domain in the proposed chloride conducting state (Chen *et al*., 2021).

### Potassium binding seals the HP2 gate

iOFS-K^+^, and iOFS-Apo imaged in the KCl and alkali-free NMDG buffers, respectively, have overall similar structures and feature well-resolved HP2s closed over the substrate-binding sites, though the hairpin conformations differ (Figure 1 and below). In iOFS-K^+^, we find a strong excess density in the substrate-binding pocket, absent in iOFS-Apo, which we interpret as a bound K^+^ ion (Figure 3A, B). The ion is coordinated by carbonyl oxygen atoms of residues in the HP2 (408, 409, and 411) and HP1 (331) tips and sidechains of D444 and T448 in TM8 (Figure 3A) and takes the place of the substrate amino group, also coordinated by D444 (Figure 3C, D) Consistently, D444S/T448A EAAT3 mutant shows impaired potassium binding and transports dicarboxylates instead of glutamate (Wang et al., 2013). Mutations of D444 to any amino acid, including glutamate, abolish transport currents, and T448A mutation alone decreases uptake by 80 % (Teichman and Kanner, 2007).

**Figure 3.**
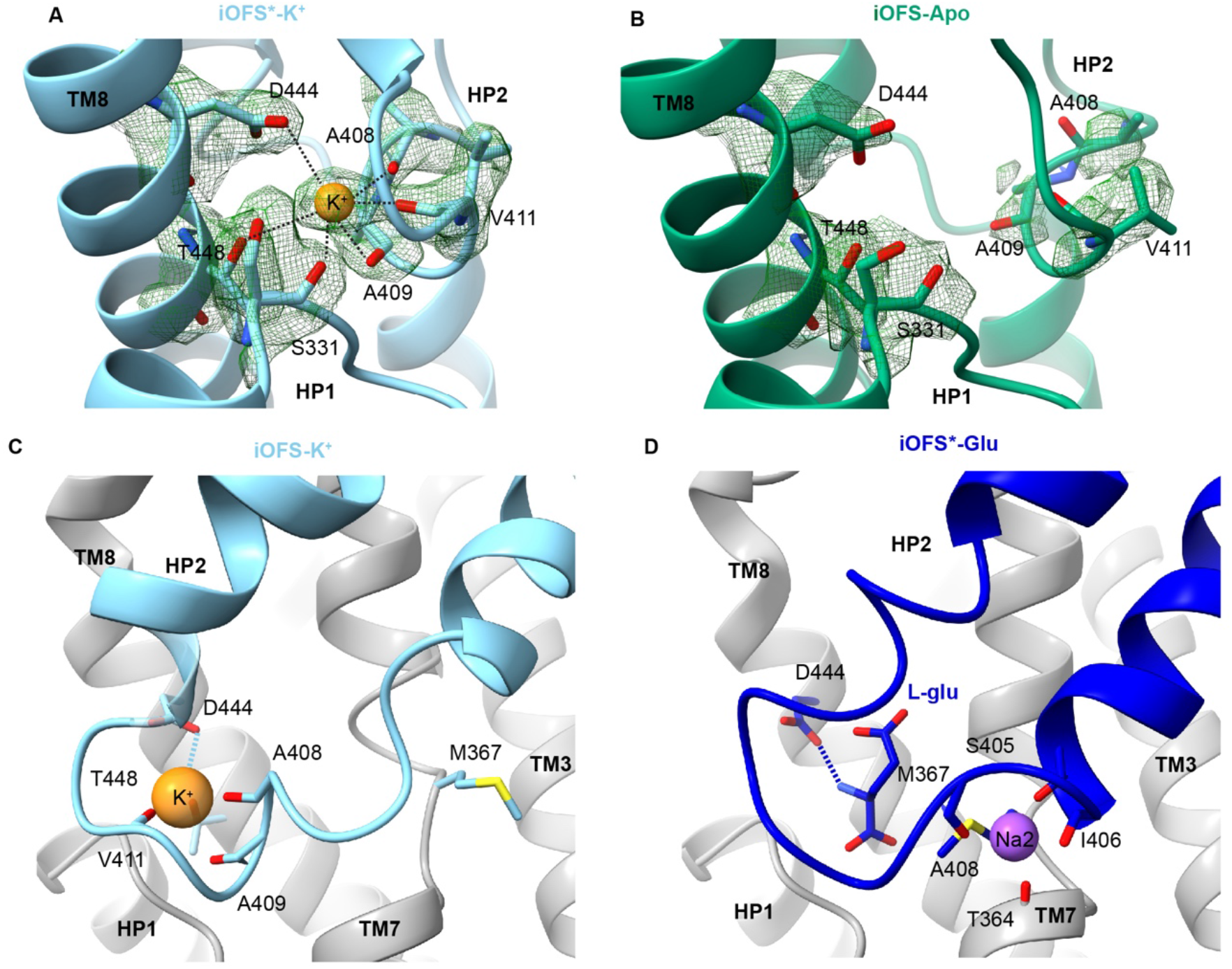
Potassium binding. (**A, B**) The substrate-binding site in iOFS-K^+^ (**A**) and iOFS-Apo (**B**). EM density, contoured at 5.5 and 5 σ, respectively, is shown as a green mesh. (**C, D**) Potassium (**C**) and glutamate (**D**) coordination. Moieties coordinating K^+^ and Na2 are emphasized as sticks. Dashed lines emphasize the shared coordination of K^+^ and L-glu by D444.

Potassium binding at this site is consistent with earlier proposals based on modeling (Holley and Kavanaugh, 2009) and the crystallographic studies of thallium binding to archaeal Glt_Ph_ (Verdon *et al*., 2014) and rubidium binding to tsEAAT1 (Canul-Tec *et al*., 2022). In tsEAAT1, rubidium bound to OFS, where HP2 remained open, and ion coordination was incomplete. Here, we observe complete octahedral ion coordination with a mean distance between the ion and the coordinating oxygens of ∼3.15 Å, within the expected range (Gagne and Hawthorne, 2016). Furthermore, the efficient occlusion of the ion by compact closed HP2 strongly supports the site as the *bona fide* potassium binding site of EAATs. Other studies proposed potassium might bind between the Na1 and Na3 sites or at the Na1 site (Kortzak et al., 2019; Wang et al., 2020). However, we see no evidence of K^+^ ions at these locations.

Strikingly, K^+^ and Na2 sites are mutually exclusive. In iOFS^*^-Glu, carbonyl oxygens of residues in the last turn of HP2a helix, 405, 406, and 408, coordinate Na2. In iOFS-K^+^, these residues unwind, while residues 408, 409, and 411 in the tip form a turn-of-a-helix-like structure, coordinating the K^+^ ion (Figure 3C, D). The overlapping binding sites for glutamate/Na2 and potassium suggest the solutes compete for binding, consistent with the antiport mechanism.

Notably, archaeal transporters do not couple to potassium counter-transport and do not bind potassium tightly (Kortzak *et al*., 2019). This is surprising because thallium bound to Apo Glt_Ph_ and K^+^ ion bound to EAAT3 share coordination by the D444 and T448. However, in archaeal transporters, glutamine replaces E374 (EAAT3 numbering), and R447, lacking its interaction partner, extends further into the substrate-binding site where it might electrostatically interfere with potassium binding. Furthermore, HP2 does not form a turn-of-a-helix structure in the archaeal transporter, and residues 409, 411, and 412 coordinate the ion in an extended loop configuration. The ability to form a helical turn in HP2 seems important for K^+^ binding because EAATs share a conserved coordinating double alanine motif AA_408-409_ with a high helical propensity. Prokaryotic homologues do not show such conservation.

### Mechanism of sodium, protons, and glutamate symport and potassium antiport

To ensure coupled transport and prevent ion gradient dissipation, iOFS-K^+^ and iOFS*-Glu should be able to translocate into IFS, but iOFS-Apo and iOFS-Na^+^ should not. What are the structural correlates distinguishing translocation-competent and -incompetent states? iOFS-Apo is structurally similar to iOFS-K^+^ (RMSD of 0.52 Å for the transport domains; Figure 4A); both feature apo configurations of the NMD and YE/DDR motifs (Figure 4B). Analogously, iOFS-Na^+^ is similar to iOFS^*^-Glu (RMSD of 0.72, Figure 4C). The NMD motif of iOFS-Na^+^ is in the bound configuration with the Na1 and Na3 sites occupied (Figure 4D). All structures have closed HP2s (Figure 1). However, in iOFS-Apo and iOFS-Na^+^, HP2 residues coordinating K^+^ and Na2 ions in iOFS-K^+^ and iOFS*-Glu, respectively, and residues 414-417 at the start of the HP2b helix unwind and form an enlarged loop with an opening in the middle (Figure 1), resembling an unzipped purse. The binding of potassium to iOFS-Apo and glutamate to iOFS-Na^+^ “zips” their HP2s, increasing their helicity and decreasing dynamics, as evidenced by the better defined HP2 EM density of iOFS-K^+^ compared to iOFS-Apo (Figure S6).

**Figure 4.**
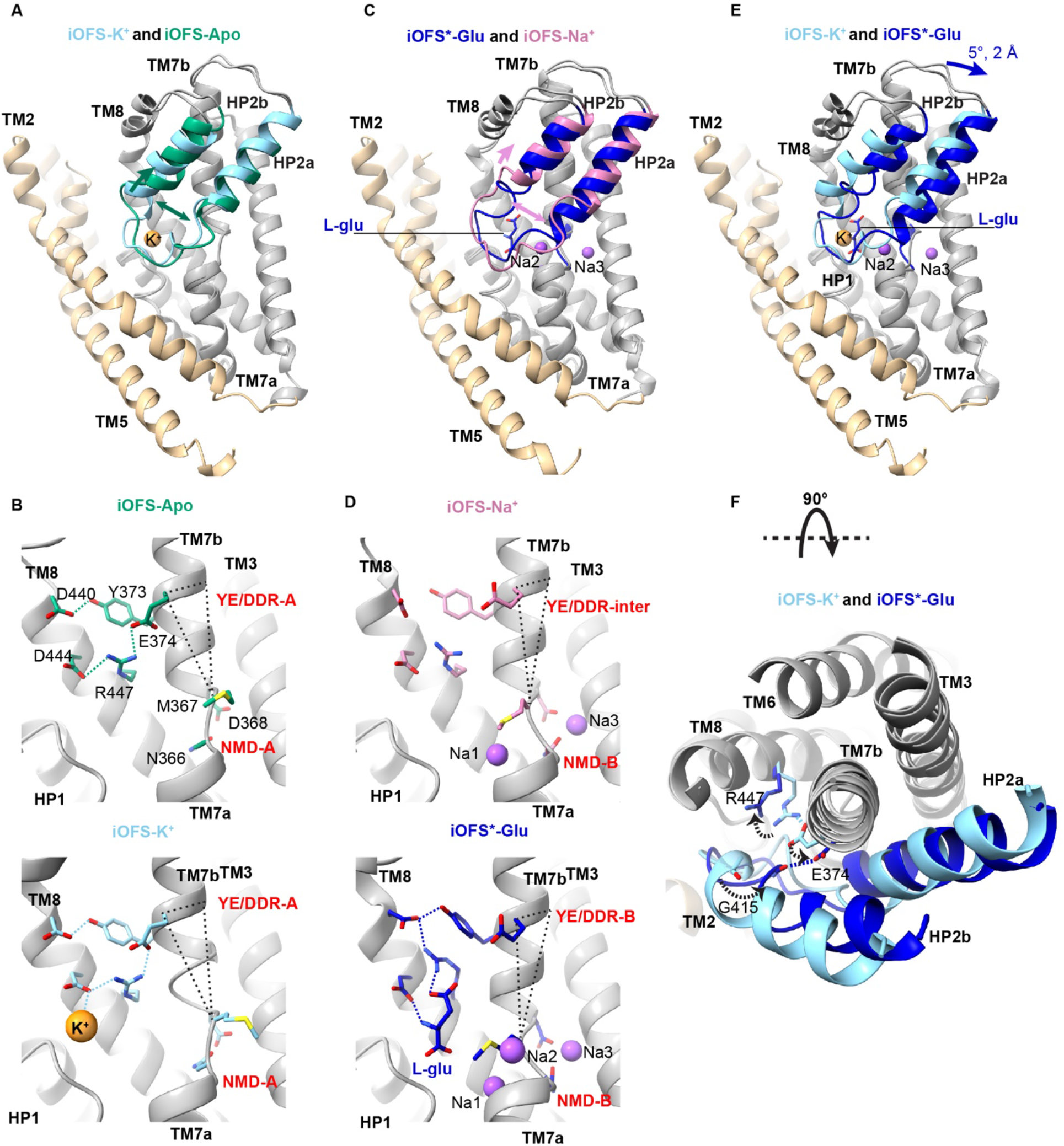
K^+^- and glutamate-mediated “zipping” of HP2. (**A, C, E**) Superimposed transport domain structures iOFS-K^+^ and iOFS-Apo (**A**), iOFS^*^-Glu and iOFS-Na^+^ (**C**), and iOFS-K^+^ and iOFS^*^-Glu (**E**). HP2 is colored light blue in iOFS-K^+^, dark green in iOFS-Apo, dark blue in iOFS^*^-Glu, and pink in iOFS-Na^+^. The protomers are superimposed on the cytoplasmic halves of the transport domain (residues 314-372 and 442-465). The scaffold domains are shown for one protomer for clarity and colored tan. Dark green, pink, and blue arrows indicate HP2 movements from iOFS-K^+^ to iOFS-Apo, iOFS^*^-Glu to iOFS-Na^+^, and iOFS-K^+^ to iOFS^*^-Glu; the K^+^ and Na^+^ ions are shown as orange and purple spheres. (**B, D**) The NMD and YE/DDR motifs in iOFS-Apo (**B**, top), iOFS-K^+^ (**B**, bottom), iOFS-Na^+^ (**D**, top), and iOFS^*^-Glu (**D**, bottom). The motifs are labeled in red and are in apo (-A), bound (-B), or intermediate (-inter) conformations. Dotted triangles connect M367, E374, and the edge of the TM7b helix. Changes in the triangle reflect rotation and tilt changes of the TM7b observed upon transition from apo to the bound conformation of the NMD motif. (**F**) Extracellular view of iOFS-K^+^ and iOFS^*^-Glu superposition. Movements of E374, G415, and R447 are shown as dotted arrows.

A comparison of EAATs to archaeal homologues, which do not couple to potassium counter-transport and can undergo elevator transition in Apo states, suggests that “zipping” HP2 is necessary for the transport domain translocation. In these transporters, HP2b remains helical even without K^+^ ions. Their YE/DDR motifs feature E374Q replacement and the glutamine hydrogen bonds to the carbonyl oxygen of G415 in HP2b, stabilizing the helix. By contrast, deprotonated E374 of the apo EAAT3 is engaged with R447 and cannot stabilize the helix, necessitating K^+^ binding to “zip” the hairpin (Figure S7A-C).

Glutamate- and K^+^-mediated “zipping” transitions differ, and HP2 shifts by 2 Å toward the cytoplasm and rotates by 5 ° in iOFS*-Glu compared to iOFS-K^+^ (Figure 4E). Sodium binding to Na1 and Na3 sites restructures the NMD motif and leads to changes in the tilt and rotation of TM7b, disengaging E374 from R447 (Figure 4D). E374 sidechain carboxyl oxygen is now within 3 Å of the carbonyl oxygen of G415 in HP2b of iOFS*-Glu (Figure 4F), suggesting the two share a proton, as previously proposed (Canul-Tec *et al*., 2022). The formed hydrogen bond likely stabilizes the HP2b helix, mimicking interactions in the archaeal transporters (Figure S7C, D). Thus, E374 protonation is coupled to glutamate-mediated HP2 “zipping”. The observations that E374Q mutation, and mutations of Y373 and R447, abolished potassium coupling and concentrative glutamate transport in EAATs (Bendahan et al., 2000; Grewer et al., 2003; Kavanaugh et al., 1997; Zhang et al., 1998) support the crucial importance of the YE/DDR motif in coupling.

iOFS-Apo should not bind and translocate glutamate to prevent the dissipation of the glutamate gradient, and iOFS-Na^+^ should not bind K^+^ ions to prevent aberrant sodium/potassium symport. Comparing iOFS-Na^+^ to iOFS-Apo structures, we observe that sodium binding leads to M367 in the NMD motif and R447 in the YE/DDR motif to swinging in and out of the binding pocket, respectively (Figure 4B, D). These transitions are necessary for R447 and M367 to coordinate glutamate and Na2, and without them, iOFS-Apo cannot bind the transmitter, as previously proposed (Arkhipova *et al*., 2020; Guskov *et al*., 2016; Qiu *et al*., 2021; Verdon *et al*., 2014; Wang and Boudker, 2020b). Furthermore, simple modeling shows that M367 must point out of the binding pocket to make space for the turn-of-the-helix in the HP2 tip coordinating K^+^ ion, thus preventing iOFS-Na^+^ from binding potassium. Conversely, M367 must point into the pocket to allow the extension of HP2a, coordinating Na2 (Figure S7E, F). Therefore, iOFS-Apo cannot bind sodium at this site.

### Protons promote OFS, gating, and sodium binding

The existence of iOFS-K^+^, iOFS-Apo, iOFS-Na^+^, and iOFS^*^-Glu raises the question: what is the role of OFS in the transport mechanism. HP2 gate is closed in all iOFS structures, suggesting that substrate and ion binding and release in these states might be kinetically slow. Moreover, in iOFS-Na^+^, R447 has already disengaged from E374 but not yet formed cation-π interaction with Y373, perhaps creating an additional kinetic barrier for glutamate binding. We hypothesize that transition into OFS facilitates efficient gating and is favored by protons, based on the following structural observations.

In contrast to iOFS-Na^+^, OFS-Na^+^ shows wide-open HP2, a fully solvent-exposed substrate-binding site (Figure 1 and Figure 5A) and R447 interacting with Y373 and in place for glutamate coordination (Figure 5B). Surprisingly, when we examined the OFS conformations imaged in the NMDG chloride and KCl buffers, we found that they closely resembled OFS-Na^+^ with wide-open HP2 gates (Figure 1 and Figure 5A). We observed no density in the K^+^-binding site, suggesting that potassium did not bind to OFS even though present in 300 mM concentration. We term these nearly identical structures (RMSD 0.57 Å) OFS-Apo_NMDG_ and OFS-Apo_K_ and refer to them collectively as OFS-Apo for short. Strikingly, the YE/DDR motifs of OFS-Apo are in the bound configuration, and their NMD motifs are in a configuration intermediate between “apo” and “bound” (Figure 5C). Inspection of the Na3 sites in OFS-Apo_NMDG_ and OFS-Apo_K_ revealed distorted geometries of the ion-coordinating residues, including shifted N366 sidechains (Figure 5C). We found excess non-protein densities in the sites, likely due to water molecules (Figure S8A). Similarly, small shifts of the coordinating N366 and N451 carbonyl oxygens distort the Na1 sites, showing no significant excess density peaks (Figure 5C and Figure S8B).

**Figure 5.**
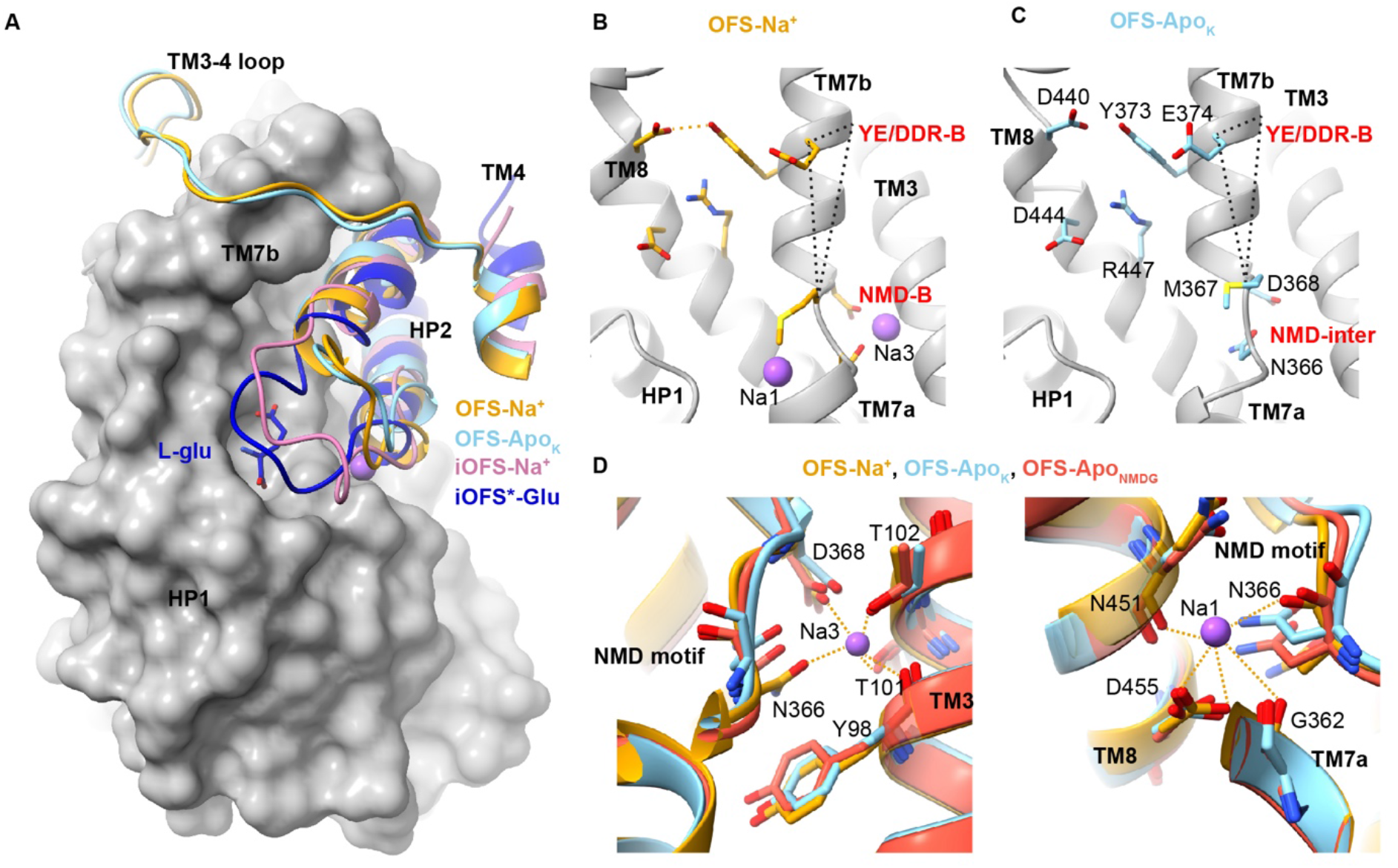
HP2 opening and the NMD and YE/DDR motifs structures in OFS. (**a**) Superposition of the transport domains of iOFS^*^-Glu (dark blue), iOFS-Na^+^ (pink), OFS-Na^+^ (orange), OFS-Apo_K_ (light blue), aligned on the cytoplasmic half of the transport domain (residues 314-372 and 442-465). The transport domain of iOFS^*^-Glu, except HP2, is shown as a gray surface. HP2 is shown in cartoon representation. TM3-4 loop and a section of TM4 are also shown for the OFS structures. (**b**,**c)** The NMD and YE/DDR motifs in OFS-Na^+^ and OFS-Apo_K_. The labeling scheme is as in Figure 4. (**d**) Comparison of the Na3 (left) and Na1 sites (right) in OFS-Na^+^ (orange), OFS-Apo_K_ (light blue), and OFS-Apo_NMDG_ (red). Orange dashed lines represent the interactions between Na^+^ ions and the coordinating atoms.

We then examined the PROPKA-predicted pKa values and solvent accessibilities of the proton carrier residue E374 in all our structures (Table S4). In iOFS^*^-Glu and OFS-Glu E374 was completely occluded from the solution and featured pKa-s of 8.4 and 8.9, respectively, suggesting the residue, paired with G415, is protonated, as expected for a fully loaded transporter. In iOFS-K^+^, E347, also buried and now paired with R447, has a pKa of 5.5 and is deprotonated, as expected for potassium counter-transport. In iOFS-Apo and iOFS-Na^+^, E374 is solvent-exposed through the “unzipped” HP2 and likely deprotonated with pKa-s of 6.5 and 7.1, respectively. In contrast, in OFS-Apo and OFS-Na^+^, E374 is buried and proximal to G415 in helical HP2b. It is likely protonated with pKa-s of 9.2, 10.0, and 7.6 for OFS-Apo_K_, OFS-Apo_NMDG_, and OFS-Na^+^, respectively.

Collectively, E374 is deprotonated in all iOFS structures, except iOFS^*^-Glu and protonated in all OFS structures. Therefore, we suggest that proton binding to E374 favors OFS with bound-like conformations of the “dancing” motifs and wide-open HP2, ready to bind sodium ions and substrate.

## Discussion

EAAT3 is a neuronal and epithelial subtype of glutamate transporters, perhaps the best functionally characterized of all EAATs (Bendahan *et al*., 2000; Furuta et al., 1997; Grewer *et al*., 2003; Grewer et al., 2000; Holmseth et al., 2012; Kanai and Hediger, 1992; Rosental et al., 2011; Teichman and Kanner, 2007; Watts et al., 2014; Watzke et al., 2001; Zerangue and Kavanaugh, 1996a; b; Zhang et al., 2007). Our eight hEAAT3 structures provide the basis for substrate recognition and selectivity and the thermodynamic coupling of substrate uptake to trans-membrane movements of three sodium ions, a proton, and a potassium ion.

EAATs transport glutamate and aspartate with similar micromolar affinities. In contrast, extensively studied archaeal transporters bind aspartate with nanomolar affinity, ten thousand times tighter than glutamate. This difference in substrate selectivity is remarkable because the substrate-binding sites are nearly identical in all structures. Moreover, hEAAT3 accommodates glutamate or aspartate upon sub-Angstrom shifts of coordinating residues and a subtle movement of the HP2b arm, regulating the pocket volume (movie S2). Recent studies on Glt_Ph_ suggest that disrupting HP2 packing against the transport domain can reduce substrate affinity (Ciftci et al., 2021; Huysmans et al., 2021). Therefore, we speculate that the tight protein packing in hyperthermophilic Glt_Ph_ accounts for its high affinity for aspartate and explains the large energetic penalty to enlarge the binding pocket for glutamate that requires movements of HP2 and altered interactions with the transport domain. In contrast, a looser packing in mammalian EAATs allows them to adapt their binding pockets to aspartate and glutamate with small energetic penalties and transport both with similar moderate affinities.

Crosslinked EAAT3-X populates OFS and occluded iOFS or iOFS^*^ (Figure 1 and Figure 6). It is possible that crosslinking stabilizes the intermediates because we observed only OFS in un-crosslinked aspartate-bound hEAAT3g (Qiu *et al*., 2021). Thus, it might be a transient state, lowly populated in the wild-type transporter. Remarkably, the state is conserved from archaea to humans (Arkhipova *et al*., 2020; Verdon and Boudker, 2012), suggesting its functional significance. iOFS conformations feature closed HP2s, while OFS-Na^+^ and OFS-Apo show the same fully open HP2 conformation, and only transmitter binding can close the gate. hEAAT3g in IFS also pictures an open HP2 (Qiu *et al*., 2021). In IFS and OFS, open HP2 interacts with the scaffold TMs 2 and 5 or the TM3-4 loop, respectively, favoring open conformations. In iOFS, the hairpins disengage from the TM3-4 loop, and their tips interact with TMs 2 and 5 as the transport domain moves inward. We suggest the discrimination between the translocation-competent and incompetent states, which is at the heart of ion-coupling mechanisms, takes place at this point. The “zipped” glutamate/3Na^+^/H^+^- or K^+^-bound transport domains can move between iOFS and IFS, while the “unzipped” domains of apo and sodium-only bound transporter cannot. This conclusion contrasts sharply with earlier hypotheses that translocation-incompetent species feature a wide-open HP2 as observed in OFS (Alleva et al., 2020; Arkhipova *et al*., 2020; Focke et al., 2011; Riederer and Valiyaveetil, 2019; Verdon *et al*., 2014; Wang and Boudker, 2020b).

**Figure 6:**
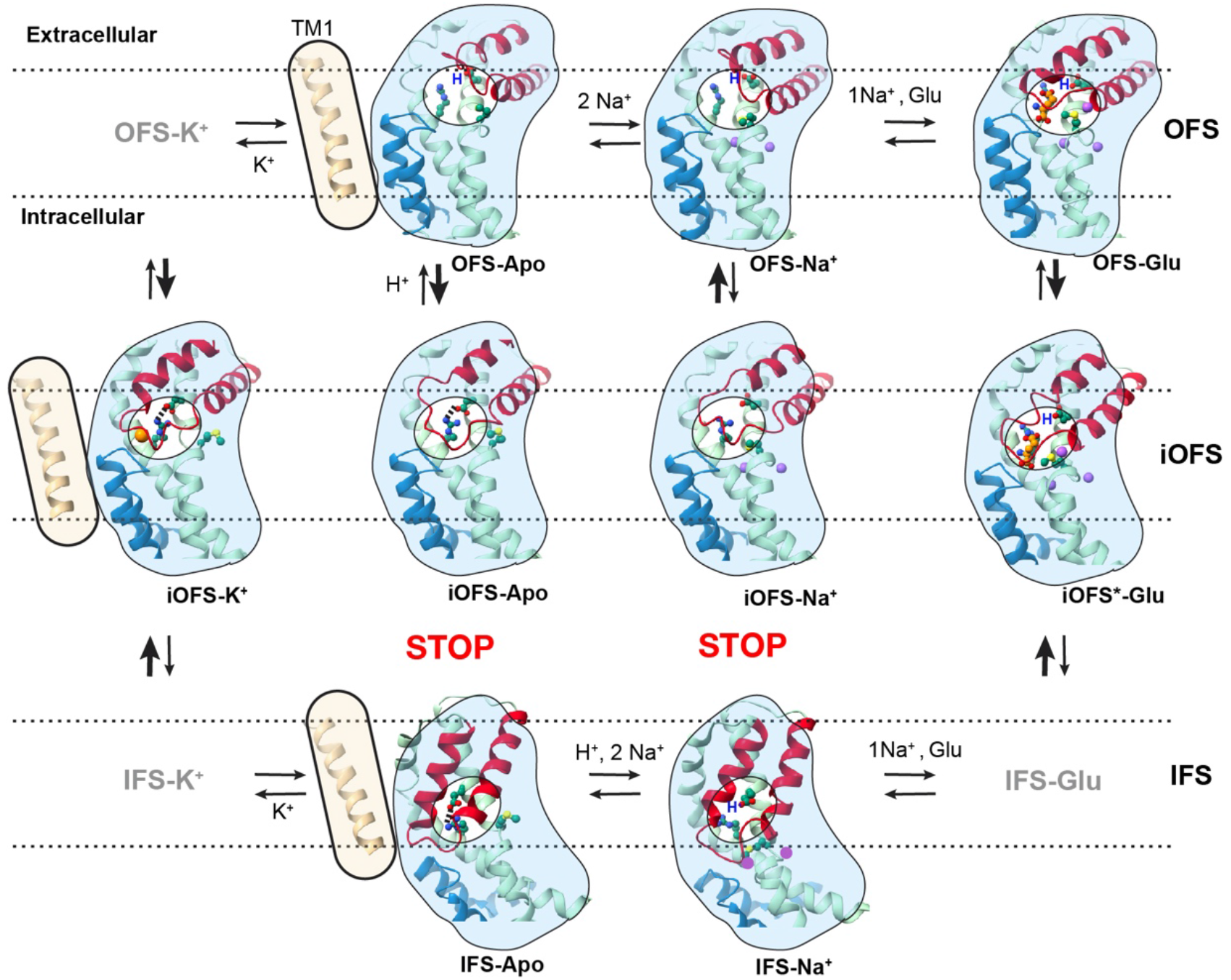
Molecular mechanism of EAAT3. Only TM1 (wheat), HP1 (dark blue), HP2 (red), and TMs 7 and 8 (light green) of a single EAAT3 protomer are shown for clarity. The substrate glutamate (orange), M367 in the NMD motif, E374, and R447 in the YE/DDR motif (dark green) are shown as ball and stick model. The K^+^ and Na^+^ ions are shown as orange and purple spheres, respectively. Blue “H” marks the proton bound to E374. The dashed lines indicate the approximate position of the lipid bilayer. The high-energy unresolved states are marked in gray. The conformational changes that are unlikely to happen are marked with red STOP sign.

The propensity of HP2 to close in iOFS explains why iOFS has the highest affinity for glutamate and potassium. Our Cryo-EM image analysis approach has previously produced state populations in good agreement with solution measurements (Huang et al., 2022). Therefore, we interpret the populations observed in 3D classifications with some confidence. We observe that iOFS and OFS populations shift from ∼7 and 93 %, respectively, when the protein is bound to sodium to 86 and 14 %, respectively, when it binds glutamate. Therefore, iOFS has ∼80 times higher affinity for glutamate than OFS. IFS binds even weaker - we observed little substrate binding in 200 mM Na^+^ and 20 mM aspartate (EMD:22024). Low affinity in IFS might ensure rapid transmitter release into the cytoplasm and fast transport rates. By contrast, slow substrate release is one of the rate-limiting steps in the tightly binding Glt_Ph_ (Ciftci *et al*., 2021). Furthermore, iOFS is the only hEAAT3 state binding K^+^ ions; neither IFS nor OFS bind even in 250 - 300 mM KCl (EMD:22023, and Figure 5). We suggest iOFS accounts for the measured 5 mM Km of potassium transport (Grewer et al., 2012). Conversely, Na^+^ ions bind ∼50-fold tighter to OFS than iOFS. We estimate that iOFS binds sodium ions with ∼1 M affinity, based on the measured sodium Km of 20 mM (Canul-Tec *et al*., 2022). Finally, the E374 pKa shifts suggest that protons bind approximately a thousand times tighter to OFS-Apo than iOFS-Apo.

Overall, we propose the following description of the transport cycle. EAAT3 OFS-Apo features an open HP2, likely protonated E374, and “bound”-like NMD and YE/DDR motifs (Figure 6). Na^+^ ions bind with high affinity to this state, and Na^+^-bound OFS might be the resting state in cells under physiologic conditions. OFS-Apo and OFS-Na^+^ can visit iOFS when transiently deprotonated but cannot progress to IFS because of “unzipped” HP2s. The glutamate and Na2 bind to OFS-Na^+^, promoting HP2 closure and transition into iOFS^*^ and IFS. Glutamate rapidly dissociates from the low-affinity IFS, followed by Na^+^ ions and the proton, yielding IFS-Na^+^ and IFS-Apo, which are open and unable to translocate back into iOFS. K^+^ ion binds in the place of the substrate leading to HP2 closure and a rapid transition into iOFS. The formation of the transient high-energy K^+^-bound occluded IFS state might be the rate-limiting step of the transport cycle, consistent with the functional data (Grewer *et al*., 2000). E374 protonation in iOFS likely triggers the transition into OFS and K^+^ release, completing the cycle.

In conclusion, we provide a complete structural description of the transport cycle of hEAAT3, highlighting the critical role of the occluded intermediate state. We also provide insights into the coupling mechanisms, energetics, substrate selectivity, and transport kinetics. It seems likely that the same structural states and coupling mechanisms exist in other glutamate transporter subtypes, though the relative state energies and, therefore, populations might vary.

## Acknowledgement

We thank Drs. Xiaoyu Wang, Krishna Reddy, and Yun Huang for the critical reading of the manuscript, data processing, and figures preparation. We thank Eric Robel for a suggestion on figures. We thank Eugene Chua at Simons EM center at New York Structural Biology Center (NYSBC), Alice Paquette, Bing Wang, and William Rice at New York University Langone’s Cryo-EM laboratory for assistance with data collection.

## Author contribution and interest conflict

B.Q. and O.B. conceived the projects; B.Q. performed all the experiments; B.Q. and O.B. analyzed data and wrote the manuscript. Competing interests: The authors declare no competing financial interests.

## Data availability

The Cryo-EM maps and atomic coordinates have been deposited in the Electron Microscopy Data Bank (EMDB) and Protein Data Bank (PDB) under accession code EMD-26985 (major conformation of EAAT3-X with 20 mM L-glu, PDB-8CTC), EMD-26986 (minor conformation of EAAT3-X with 20 mM L-glu, PDB-8CTD), EMD-26997 (major conformation of EAAT3-X with 300 mM KCl, PDB-8CUA), EMD-26998 (minor conformation of EAAT3-X with 300 mM KCl, PDB-8CUD), EMD-27000 (major conformation of EAAT3-X with 150 mM NMDG-Cl, PDB-8CUI), EMD-27001 (minor conformation of EAAT3-X with 150 mM NMDG-Cl, PDB-8CUJ), EMD-27006 (major conformation of EAAT3-X with 300 mM NaCl, PDB-8CV2), and EMD-27007 (minor conformation of EAAT3-X with 300 mM NaCl, PDB-8CV3).

## Materials and Methods

### Protein expression and purification

The codon-optimized full-length human EAAT3 with N178T and N195T mutations (hEAAT3g) (Qiu *et al*., 2021) was cloned into a modified pCDNA 3.1 (*Invitrogen*) plasmid which has N-terminal Strep-II tag followed by GFP and PreScission protease site. Cysteine residues C9, C100, C158, and C219 were mutated to alanines and C256 to tryptophan for yielding MiniCys EAAT3 construct. Double cysteine mutation K269C/W441C was introduced in MiniCys EAAT3 for crosslinking in OFS. All mutations were prepared by site-directed mutagenesis (*Agilent*). Transient transfections and protein purifications were performed as described previously (Qiu *et al*., 2021).

Briefly, 1 litter human embryonic kidney (HEK) 293F cell cultured in Freestyle 293 medium (*Gibico*) with the density of about 2.5 million cell/ml were transfected using 3 mg plasmid by polyethylenimine (PEI, *Polysciences*) with a 1:3 plasmid to PEI weight ratio. Transfected cells were diluted with 1 litter fresh Freestyle 293 medium 6 hours after transfection, and then 2.2 mM valproic acid sodium (*Sigma-Aldrich*) was added to cells to boost protein expression 12 hours later. About 48 hours after transfection, the cells were collected by centrifugation at 4000g for 10 min at 4°C. The cells were resuspended in lysis buffer containing 50 mM Tris-Cl (pH 8.0), 1 mM L-asp, 1 mM EDTA, 1 mM tris(2-carboxyethyl) phosphine (TCEP), 1 mM phenylmethylsulfonyl fluoride (PMSF), and 1:200 dilution of protease inhibitor cocktail (P8340, *Sigma-Aldrich*) and disrupted with an EmulsiFlex-C3 cell homogenizer (*Avestin*). The cell debris was removed by centrifugation at 10,000g for 15 min at 4°C, and membrane pellets were collected by ultracentrifugation at 186,000g for 1 hour at 4°C. The membrane pellets were homogenized in lysis buffer supplemented with 200 mM NaCl and 10% (v/v) glycerol (solubilization buffer) and then incubated with 1% dodecyl-β-D-maltopyranoside (DDM; *Anatrace*) 0.2% cholesteryl hemisuccinate (CHS; *Sigma-Aldrich*) at 4°C overnight. Insoluble material was removed by ultracentrifugation at 186,000g for 1 hour at 4°C, and the supernatant was incubated with Strep-Tactin Sepharose resin (*GE Healthcare)* for 1 hour at 4°C. The resin was washed by eight column volumes of the wash buffer containing 50 mM Tris-Cl (pH 8.0), 200 mM NaCl, 0.06% glyco-diosgenin (GDN; *Anatrace*), 1 mM TCEP, 5% glycerol and 1 mM L-asp. Protein was eluted with four column volumes of the wash buffer supplemented with 2.5 mM D-desthiobiotin (elution buffer). The N-terminal Strep II and eGFP tags were cleaved by incubating the protein with homemade PreScission protease at a 40:1 protein to protease ratio overnight at 4°C. Eluted protein was further purified by size exclusion chromatography (SEC) using Superose 6 Increase 10/300 column (*GE Healthcare*), pre-equilibrated with the desired buffers.

To prepare crosslinked EAAT3-X with 20 mM L-glu, the protein was first purified by SEC in a buffer containing 20 mM Hepes-Tris pH 7.4, 200 mM NaCl, 0.01% GDN, and 1 mM L-asp. Peak SEC fractions were pooled, concentrated to ∼0.5 mg/ml, and incubated with HgCl_2_ at a 1:20 protein to Hg^2+^ molar ratio for 15 min at room temperature. Crosslinked EAAT3-X was further exchanged into a buffer containing no sodium or aspartate by SEC (20 mM Hepes-Tris pH 7.4, 100 mM choline chloride, 0.01% GDN). The peak fractions were supplemented with L-glutamate by diluting ∼1,000-fold into a buffer containing 20 mM Hepes-Tris pH 7.4, 200 mM NaCl, 20 mM L-glutamate, 0.01% GDN, and concentrated using 100 kD MWCO concentrators (*Amicon*). EAAT3-X protein samples in 300 mM KCl and 150 mM N-Methyl-D-glucamine (NMDG) chloride were prepared by SEC in buffers containing 20 mM Hepes-Tris pH 7.4, 0.01% GDN, and 300 mM KCl or 150 mM NMDG chloride. To prepare the EAAT3-X sample in 300 mM NaCl, protein in 150 mM NMDG buffer was diluted ∼1,000-fold into a buffer containing 20 mM Hepes-Tris pH 7.4, 300 mM NaCl and 0.01% GDN. Notably, we prepared NMDG and Na^+^ samples using the same purified crosslinked protein.

### Cry-EM data acquisition

3.5 μl of protein samples at 4-6 mg/ml were applied to glow-discharged QF R1.2/1.3 300 mesh gold grids (*Quantifoil*, Großlöbichau, Germany). Grids were blotted for 3 s at 4°C and 100% humidity and plunge-frozen into liquid ethane using FEI Mark IV Vitrobot (*FEI*, part of *Thermo Fisher Scientific*, Hillsboro, OR). Dataset for EAAT3-X with 20 mM L-glutamate was collected using Leginon (Suloway et al., 2005) at New York Structural Biology Center (NYSBC). Datasets for crosslinked EAAT3-X with 300 mM KCl, 150mM NMDG, and 300 mM NaCl were collected using Leginon (Suloway *et al*., 2005) at NYU Langone Health’s Cryo-Electron Microscopy Laboratory. Detailed parameters of cryo-electron microscopy data collection were listed in Supplementary Table 1.

### Image processing

All the datasets were processed using a combination of Relion 3.1.0 (Scheres, 2012) and CryoSPARC (Punjani et al., 2017) programs. Movie drift correction was performed using Motioncor2 (Zheng et al., 2017), and CTF parameters of the micrographs were estimated using CTFFIND (Rohou and Grigorieff, 2015) in Relion 3.1.0.

For the EAAT3-X in the 20 mM L-glu dataset, 8,511,485 particles were auto-picked from 6,361 micrographs by Laplacian-of-Gaussian (LoG). The particles were extracted using a box size of 128 pixels with 2x binning and imported into CryoSPARC for 2D classification. 1,087,381 particles showing secondary features were selected and subjected to 3 consecutive rounds of *ab initio* reconstruction. 88,094 particles producing a volume with most features were refined to 6.2 Å by non-uniform refinement (Punjani et al., 2020) (hereafter, NUR), applying C1 symmetry. Next, 6,867,502 particles after 2D cleaning were subjected to a round of heterogeneous refinement using one good volume from NUR and 8 decoy noise volumes generated by running 1-2 iterations of *ab initio* reconstruction. This procedure, termed heterogeneous refinement for cleaning (HRC), sorts informative particles from noise. 1,726,968 particles were selected and refined to 6.3 Å by NUR in C1. The particles were re-imported into Relion using PyEM and, after unbinning, re-extracted using the box size of 256 pixels. These particles were re-imported to CryoSPARC and subjected to a round of HRC. The selected 682, 971 particles were refined to 3.35 Å by NUR with C3 symmetry. The refined particles went through two rounds of polishing in Relion, HRC, and NUC steps. The final map was refined to 2.8 Å with C3 symmetry using 496, 972 particles. Symmetry expansion and local classification with a mask applied over the transport domain yielded 14 % of protomers in a different conformation. These particles yielded a 3.42 Å resolution map after local refinement with a mask applied over the whole protomer and postprocessing in Relion.

For the EAAT3-X in 300 mM KCl dataset, 3,502,663 unbinned particles extracted with a 300-pixel box size from 5,116 micrographs were imported into CryoSPARC for 2D classification. 892,591 particles showing secondary features were selected, and then a 3.35 Å model was generated using 154,013 particles after 2 rounds of *ab initio* and 1 round of NUR in C1 symmetry. 3 million particles selected after 2D cleaning were subjected to 2 rounds of HRC, using one good volume from NUR (3.35 Å) and 7 decoy noise *ab initio* volumes. 426,862 selected particles were refined to 2.82 Å by NUC with C3 symmetry. Polishing in Relion, HRC, and NUC steps were repeated twice. Finally, 404,729 particles were refined to 2.44 Å by NUC with C3 symmetry. Symmetry expansion and local classification identified ∼10% of protomers in a different conformation. The particles from the minor class yielded a 2.93 Å map after local refinement and postprocessing in Relion. Figure S4 shows the image processing flowchart for the dataset. The image processing of EAAT3-X with 150 NMDG and 300 mM NaCl datasets was similar to the KCl dataset. The data processing information is in Supplementary Tables 2 and 3.

### Model building and refinement

For the model building, structures of hEAAT3g (PDB accession codes 6×2l, 6×3f, and 6×2z) were first fitted into maps using ChimeraX (Goddard et al., 2018). The models were manually adjusted in COOT (Emsley et al., 2010), and subjected to real space refinement in Phenix (Adams et al., 2010). The cross-validation was performed by displacing atoms in the final model by 0.3 Å, refining the displaced model against the first unfiltered half-map (FSC-work). FSC curves were then calculated between the FSC-work model and the second unfiltered half-map (FSC-free) and between the refined model and the full map (FSC-sum). The structure figures were prepared by ChimeraX and Pymol (DeLano Scientific). The tunnel through protein structure was calculated using CARVER3.0 (Chovancova et al., 2012). The pKa of E374 was calculated by PROPKA 3.0 (Olsson et al., 2011). The solvent accessible surface area of E374 was calculated by PISA (Krissinel and Henrick, 2007).

### Proteoliposome reconstitution and solid supported membrane (SSM) assay

The proteoliposome reconstitution and SSM assay were performed as before described (Qiu *et al*., 2021). Liposomes were prepared using 1-palmitoyl-2-oleoyl-sn-glycero-3-phosphocholine (POPC, *Avanti Polar Lipids*), 1-palmitoyl-2-oleoyl-sn-glycero-3-phosphoethanolamine (POPE, *Avanti Polar Lipids*) and CHS at a 5:5:2 ratio. The lipids in chloroform were dried and rehydrated at 20 mg/ml by 10 free-thaw cycles in 50mM HEPES-Tris buffer, pH 7.4, forming multilamellar liposomes. The liposomes were diluted to 4 mg/ml in a buffer containing 50 mM Hepes/NaOH, pH7.4, 200 mM NaCl, 1mM TCEP, 1mM L-asp, and extruded 11 times through 400 nm polycarbonate membranes (*Avanti Polar Lipids*) by a syringe extruder (*Avanti Polar Lipids*). Then the unilamellar liposomes were destabilized with DDM at a 1:0.75 lipid to detergent ratio at 23°C for 15 min. Purified MinCys EAAT3 and MinCys EAAT3 C343V were added to the liposomes and incubated for 30 min at 23°C at a 1:10 protein to lipid ratio. Detergents were removed by incubation with 100 mg/ml Bio-Beads SM-2 (*Bio-Rad*) for 1h at 23°C, 1h at 4°C (3 times), overnight at 4°C, and 1h at 4°C. The proteoliposomes were collected by centrifugation at 40,000 rpm for 45 min at 4°C, and then diluted into buffer containing 100 mM potassium phosphate, pH 7.4, 2 mM MgSO_4_ (SSM assay resting buffer). The re-suspended was subjected to freeze-thaw in liquid nitrogen. The centrifugation and freeze-thaw steps were repeated 3 times for complete buffer exchange. The SSM assays were performed on a SURFE2R N1 instrument (*Nanion Technologies*). Briefly, proteoliposomes from last step were extruded 11 times through 400 nm polycarbonate membranes and coated onto SF-N1 sensor (*Nanion Technologies*) according to the instrument manual. The non-activating buffer containing 100 mM sodium phosphate, pH 7.4, and 2 mM MgSO_4_ flowed through the sensor at 200 μl/s flow rate to build the ionic gradient against the coated proteoliposomes. The transport-coupled current was generated by flowing activating buffer containing 100 mM sodium phosphate, pH 7.4, 2 mM MgSO_4_ and 3mM L-asp. Finally, the sensor was rinsed in the resting buffer to restore the sensor. At least two sensors for each proteoliposome preparation were recorded, producing similar results.

