## Supplementary_Material for "Dancing with the ions: symport and antiport mechanisms of human glutamate transporters"

**This PDF file includes:**

Figure S1-8

Table S1-4

Legends of Movie S1, 2

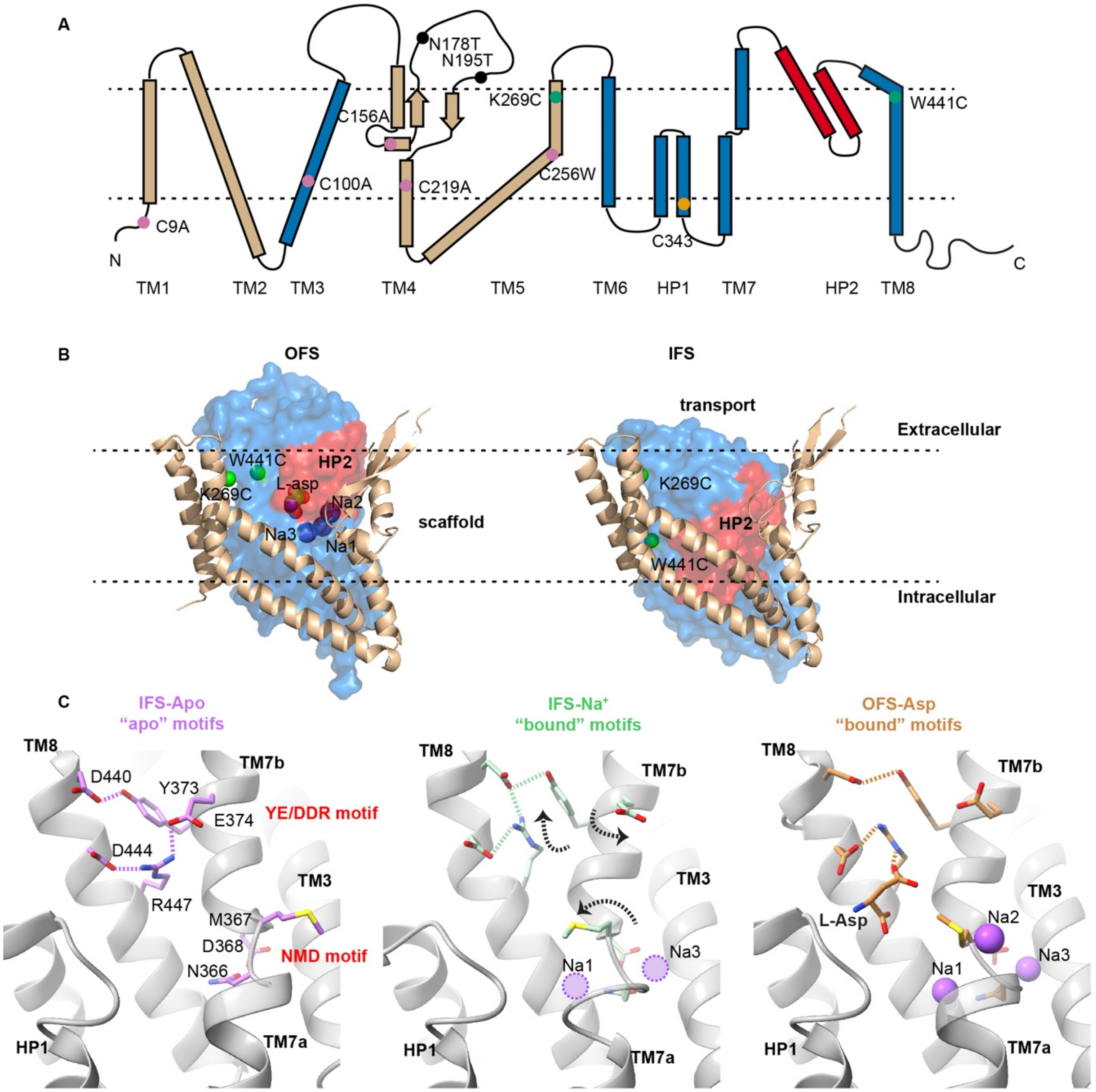

**Figure S1: EAAT3 topology, elevator transport mechanism, and "dancing" motifs.** (**A**) The EAAT3 topology with scaffold domain colored wheat, transport domain blue, and HP2 red. The pink circles represent cysteines mutated to alanines or a tryptophan; the orange circle corresponds to the highly conserved cysteine in HP1; the green circles represent K269 and W441 mutated to cysteine for crosslinking. (**B**) The elevator movements of the transport domain. Shown are single hEAAT3g protomers in the outward-facing aspartate-bound state (OFS, left, PDB ID 6x2z) and inward-facing sodium-bound state (IFS, right, PDB ID 6x2l). The scaffold and transport domains are in a cartoon and surface representation, respectively. Green spheres emphasize the K269 and W441 Cα-s. The bound aspartate and sodium ions are shown as spheres and colored by atom type. (**C**) The "dancing" NMD and YE/DDR motifs form the substrate- and ion-binding sites in IFS-Apo (left, PDB ID 6x3f), IFS-$Na^+$ (middle, PDB ID 6x2l), and OFS-Asp (right, PDB ID 6x2z) states. HP2, occluding the sites, is removed for clarity. Dashed lines mark key interactions in the motifs, which transition from an “apo” to a “bound” configuration upon sodium binding, prepping the sites for the consequent transmitter binding. Dotted circles represent bound unresolved $Na^+$ ions in IFS-$Na^+$.

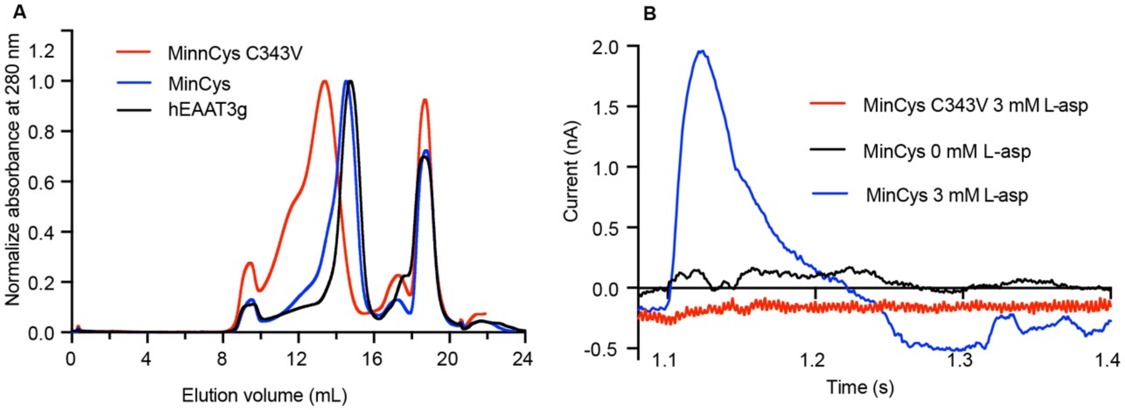

**Figure S2: Characterization of MinCys EAAT3.** (**A**) SEC elution profiles of hEAAT3g (black), MinCys (blue), and MinCys C343V mutant (red). The absorbance at 280 nm was normalized by the peak maximum. (**B**) MinCys displays substrate transport currents (blue) in solid supported membrane (SSM) assays. In contrast, C343V mutant (red) is indistinguishable from the control (black). Other C343 mutants showed similar unfavorable behaviors.

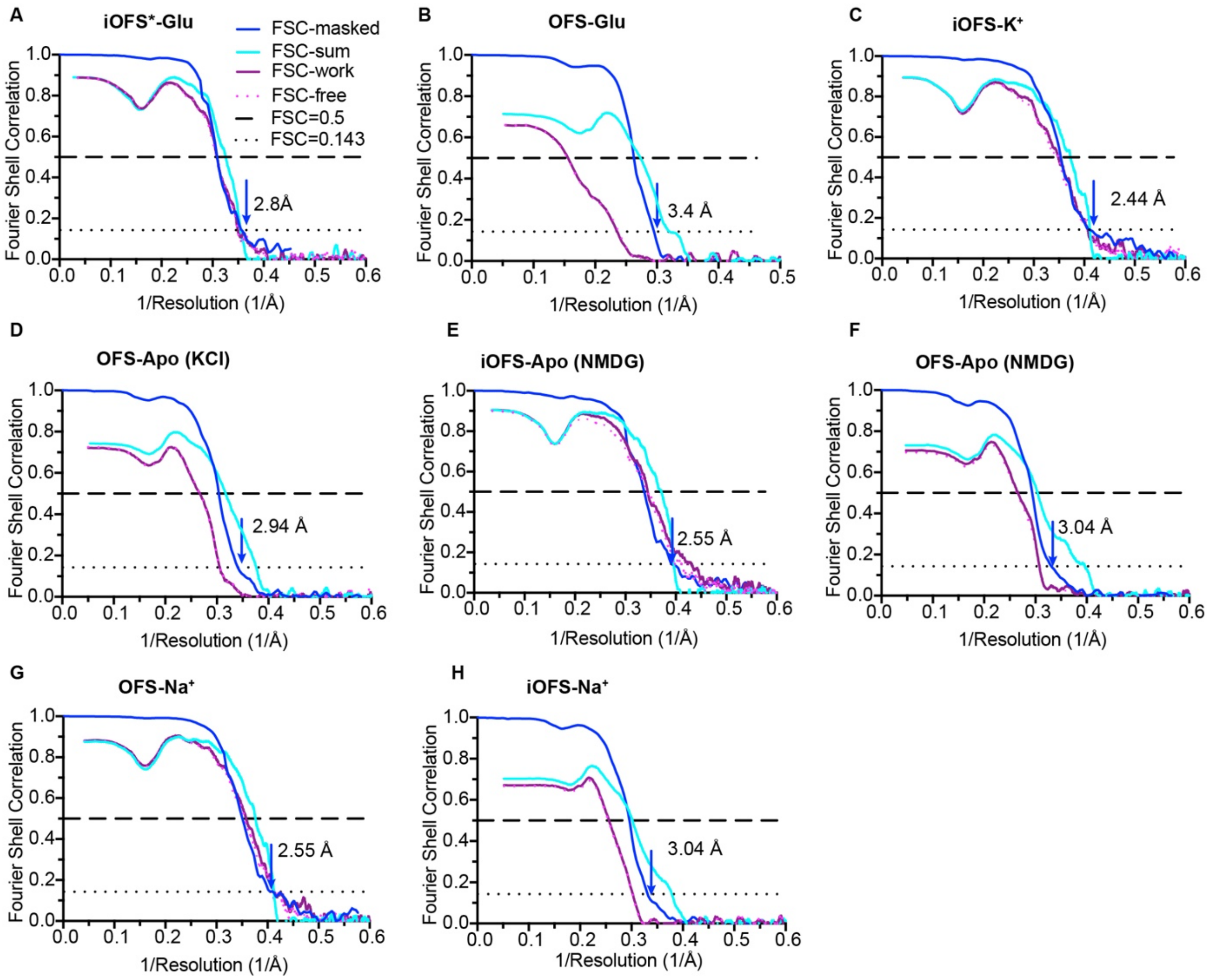

**Figure S3: Maps and models validation**. Fourier Shell Correlation (FSC) curves for the density maps, and maps and models validations for EAAT3-X in the following states: iOFS*-Glu (**A**), OFS-Glu (**B**), iOFS-$K^+$ (**B**), OFS-Apo in KCl (**D**), iOFS-Apo in NMDG (**E**), OFS-Apo in NMDG (**F**), OFS-$Na^+$ (**G**), and iOFS-$Na^+$ (**H**). Shown are the FSC curves for the density maps (blue) and the FSC curves for the refined models versus full maps (cyan) and half maps for cross-validation (purple and pink dots). Dashed and dotted lines correspond to the FSC values of 0.5 and 0.143, respectively.

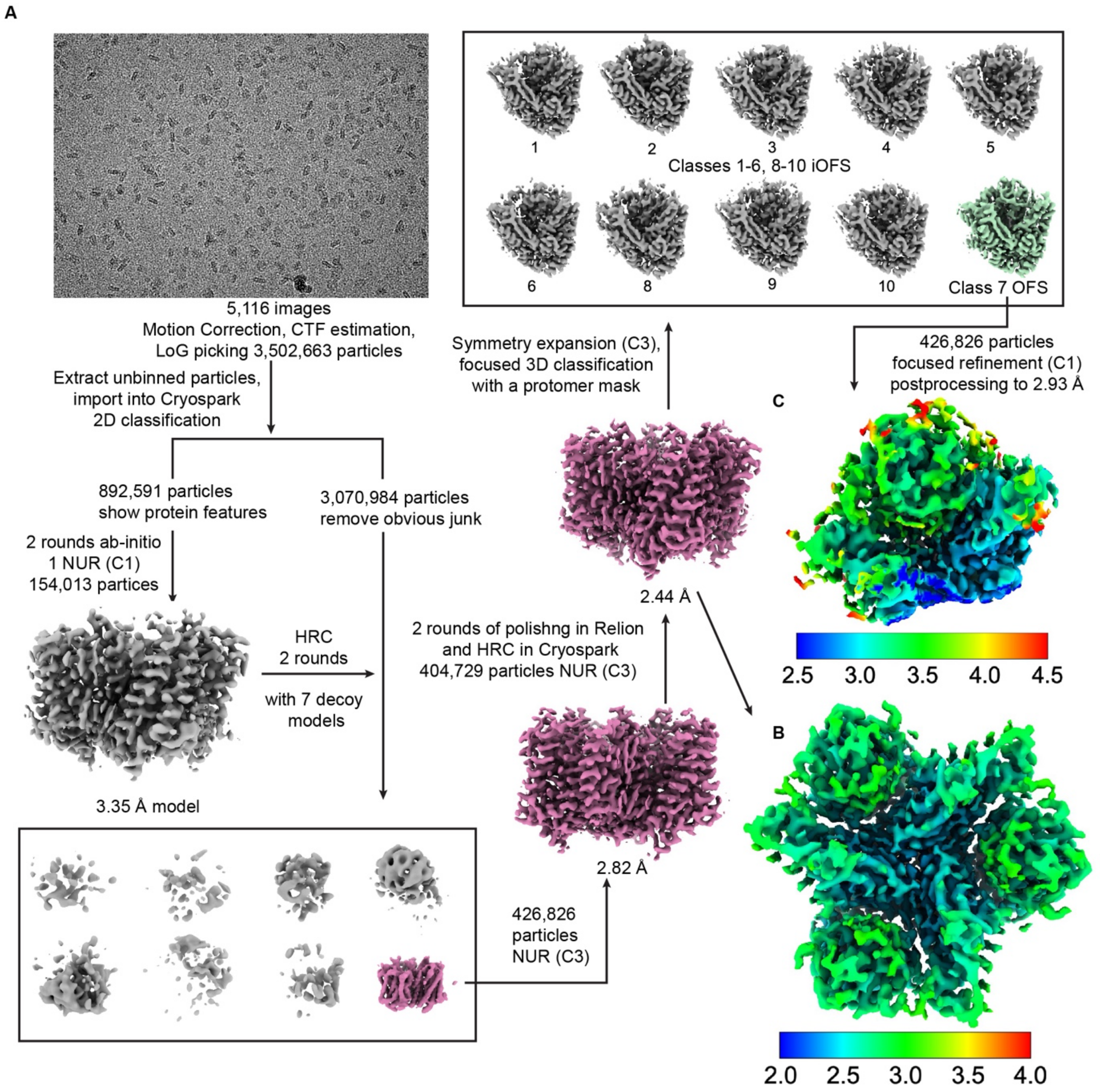

**Figure S4: Example of the data processing flowchart for EAAT3-X in 300 mM KCl.** (**A**), An example of an aligned image and the data processing flowchart. (**b**, **c**), Local resolution distribution of iOFS-K map refined in C3 (**B**) and the minor OFS-Apo protomer, showing no density for a bound $K^+$ ion (**C**).

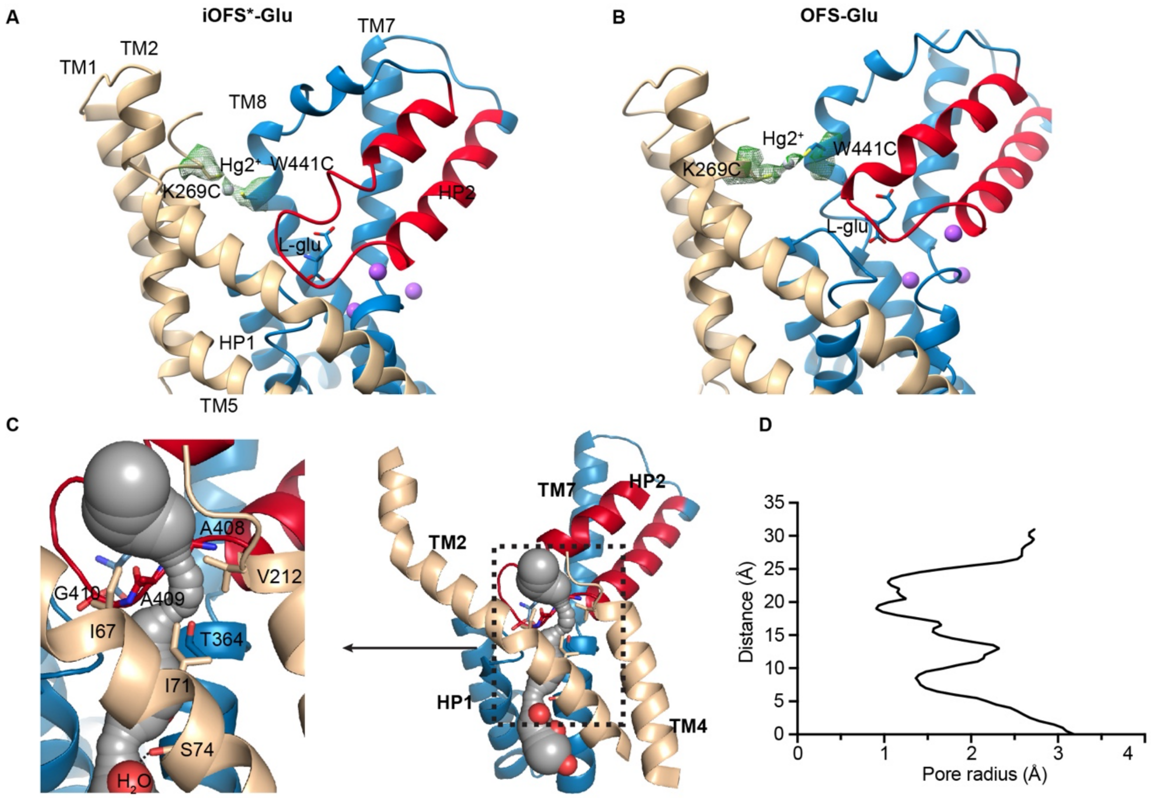

**Figure S5: Glutamate-bound EAAT3-X in iOFS* and OFS.** A cross-link between K269C and W441C residues in iOFS*-Glu (**A**) and OFS-Glu (**B**). Shown are single protomers in cartoon representation colored as in Supplementary Figure 1 with TMs 3, 4, and 6 removed for clarity. Green mesh is the EM density around K269C, W441C, and $Hg^{2+}$ ions, contoured at 5 σ. (**C**), The tunnel in iOFS*-Glu is calculated by CAVER 3.0 and shown as gray spheres. The observed solvent molecules (red spheres) are visible in the lower part of the tunnel. Residues forming the extracellular constriction and S74 implicated in anion selectivity are shown as sticks. (**D)**, The pore radius along the tunnel. Zero on the Y axis is set to the location of the second water in the cytoplasmic vestibule.

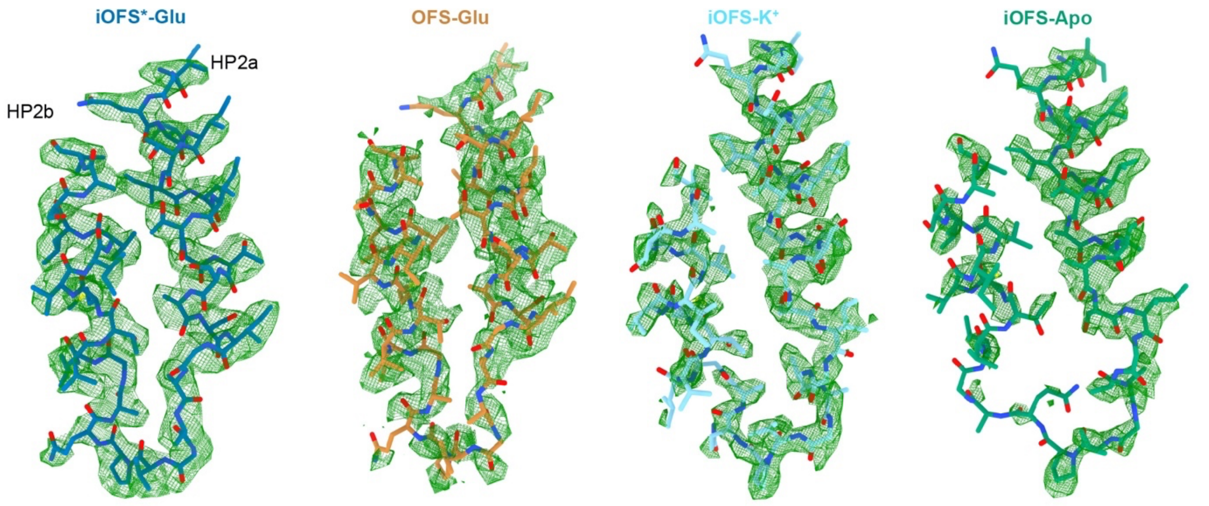

**Figure S6: HP2 EM density maps.** The maps, shown as green mesh, are contoured at 5σ around HP2 of, from left to right, iOFS$^{*}$-Glu, OFS-Glu, iOFS-K$^{+}$, and iOFS-Apo states. Modeled HP2 is shown in stick representation.

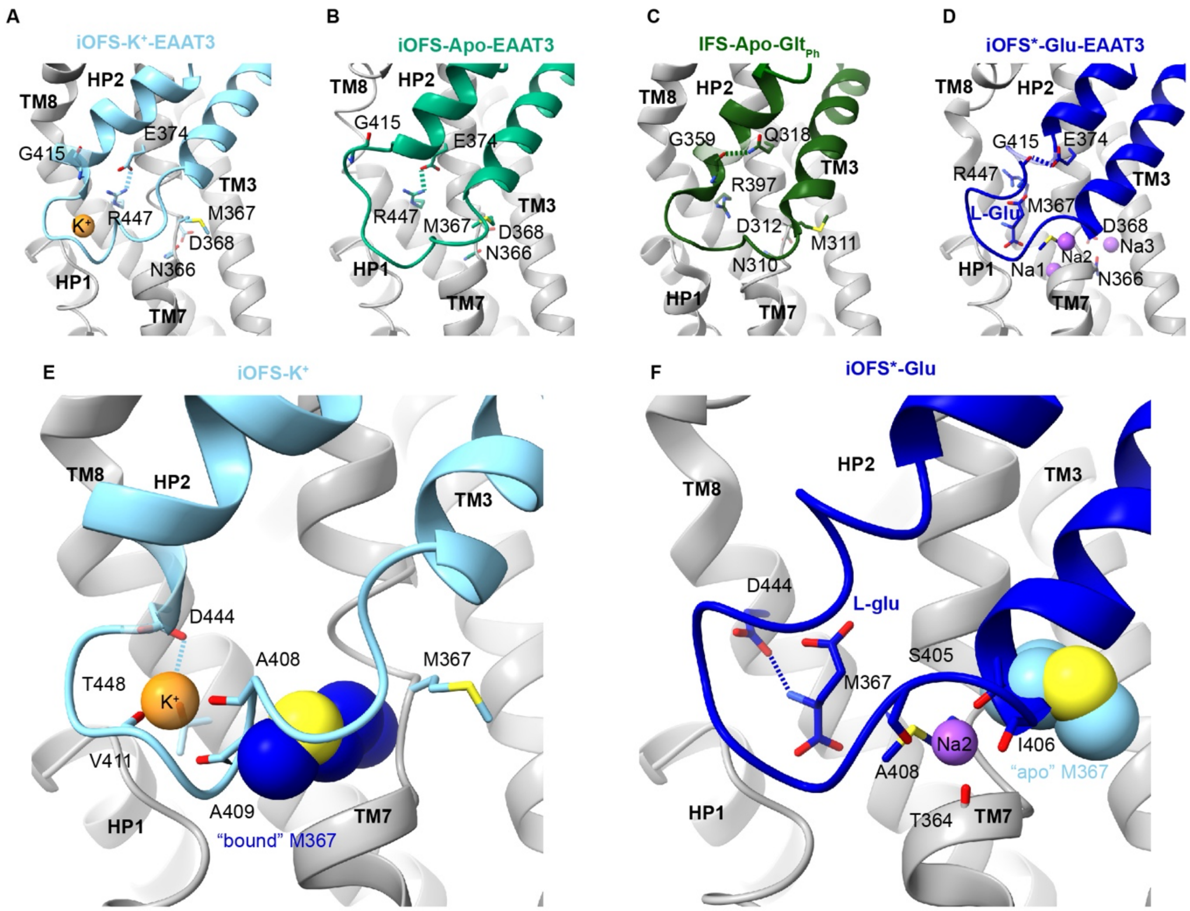

**Figure S7: The NMD and YE/DDR motifs mediate ions coupling in EAAT3.** (**A-D**), The YE/DDR motifs in EAAT3 iOFS-$K^+$, EAAT3 iOFS-Apo, $Glt_{Ph}$-Apo, and EAAT3 $iOFS^*$-Glu. Structures are superposed on the cytoplasmic halves of their transport domains (residues 314-372 and 442-465 for EAAT and residues 259-317, 392-415 for $Glt_{Ph}$, PDB ID 4oye). The interactions between E374 and R447 or G415 in EAAT3 and between Q374 and G415 in $Glt_{Ph}$ are shown as dashed lines. (**E**, **F**) Orientation of M367 in the NMD motif determines whether HP2 can coordinate $K^+$ ion or Glu/Na2. The HP2 in the potassium-bound conformation (**E**) would sterically clash with the M367 in glutamate-bound conformation, shown as dark blue spheres. Conversely, HP2 in the glutamate-bound conformation (**F**) would clash with M367 in the apo conformation (light blue spheres).

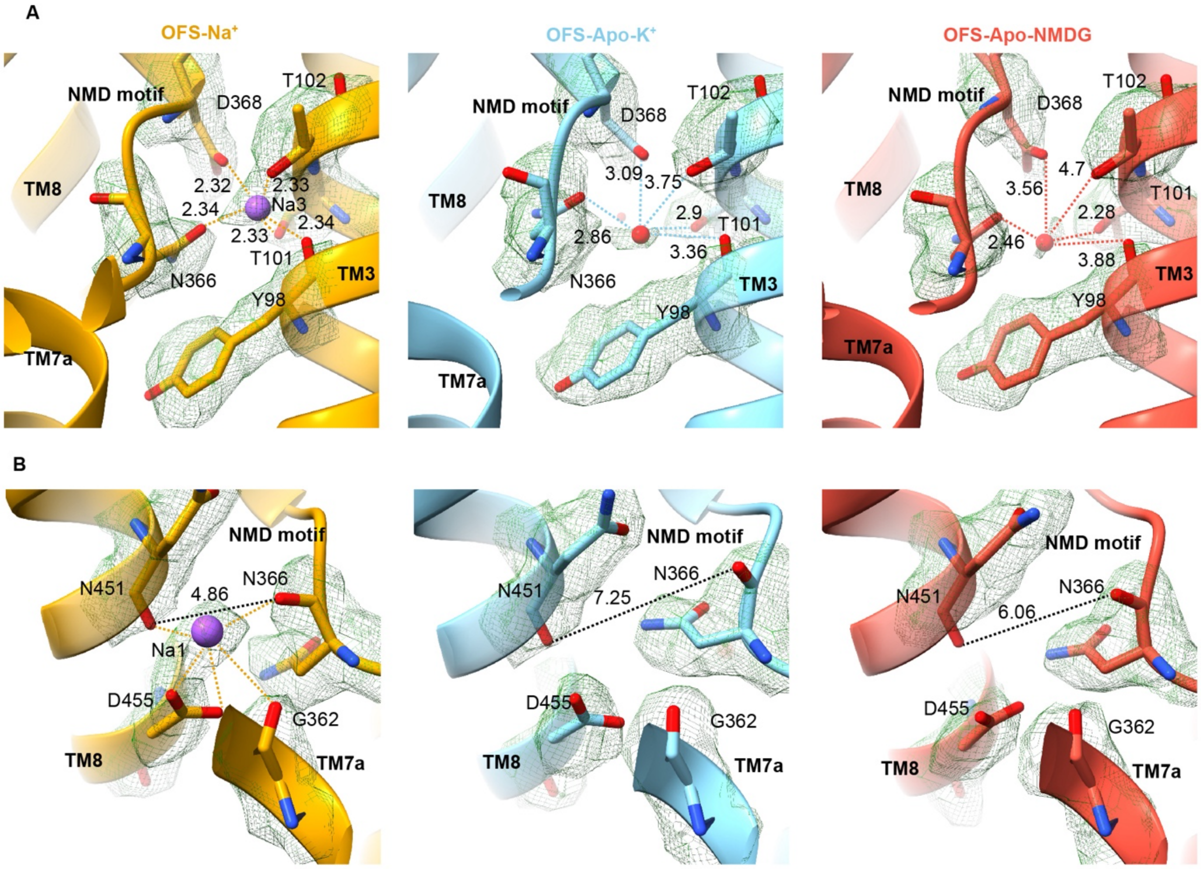

**Figure S8: Distorted sodium sites in OFS-Apo.** EM density and geometry of Na3 (**A**) and Na1 (**B**) sites of OFS-Na$^+$ (left), OFS-Apo$_K$ in 300 mM KCl (middle), and OFS-Apo$_{NMDG}$ 150 mM NMDG chloride (right). The density maps, shown as green mesh, are contoured at 5.5, 5, and 5 σ, respectively. The dashed yellow lines in OFS-Na$^+$ emphasize interactions between the Na$^+$ ions and coordinating oxygens. The distances between the Na$^+$ ion or water in the Na3 site and the coordinating oxygens are shown in (**A**). In OFS-Apo structures, a solvent molecule replaces the Na$^+$ ion in the Na3 site, and the distances to coordinating moieties increase (**A**). The dashed black lines in (**B**) mark the distances between the carbonyl oxygens of N366 and N451. The distance increases observed in KCl and NMDG chloride reflect the distortions of the Na1 site in OFS-Apo.

**Table S1: Cryo-EM data collection**

| | EAAT3-X | | | |
|---|---|---|---|---|
| | 20 mM L-glu | 300 mM KCl | 150 mM NMDG-Cl | 300 mM NaCl |
| Microscope/camera | Krios/K3 | Krios/K3 | Krios/K3 | Krios/K3 |
| Voltage (kV) | 300 | 300 | 300 | 300 |
| Energy filter | 20 eV | 20 eV | 20 eV | 20 eV |
| Magnification | 81,000 X | 105,000 X | 105,000 X | 105,000 X |
| Super-resolution pixel size (Å) | 0.5413 | 0.426 | 0.426 | 0.426 |
| Dose (e-/Å2) | 50.27 | 51.10 | 50.73 | 57.52 |
| Number of frames | 40 | 48 | 52 | 40 |
| Exposure time (s) | 2 | 2.4 | 2.6 | 2.4 |
| Defocus range (μm) | -0.8 ~ -2.5 | -1.3 ~ -2.0 | -1.3 ~ -1.6 | -1.0 ~ -1.8 |

**Table S2: Cryo-EM data refinement and validation of the major conformation in C3**

| | EAAT3-X 20 mM L-glu<br>EMD-26985<br>PDB-8CTC | EAAT3-X 300 mM KCl<br>EMD-26997<br>PDB-8CUA | EAAT3-X 150 mM NMDG-Cl<br>EMD-27000<br>PDB-8CUI | EAAT3-X 300 mM NaCl<br>EMD-27006<br>PDB-8CV2 |
|---|---|---|---|---|
| **Data collection and processing** | | | | |
| Symmetry imposed | C3 | C3 | C3 | C3 |
| Initial particle images (no.) | 8,511,485 | 3,502,663 | 2,553,613 | 2,949,270 |
| Final particle images (no.) | 496,972 | 404,729 | 210,303 | 519,857 |
| Map resolution (Å) | 2.80 | 2.44 | 2.55 | 2.44 |
| FSC threshold | 0.143 | 0.143 | 0.143 | 0.143 |
| Map resolution range (Å) | 2.46 ~ 39.28 | 1.86 ~ 39.27 | 2.23 ~ 39.19 | 2.18 ~ 37.14 |
| **Refinement** | | | | |
| Initial model used (PDB code) | 6x2z, 6x2l | 6x2z, 6x3e | 6x2z, 6x3e | 6x2z, 6x2l |
| Model resolution (Å) | 3.07 | 2.68 | 2.69 | 2.66 |
| FSC threshold | 0.5 | 0.5 | 0.5 | 0.5 |
| Model resolution range (Å) | 3.07 ~ 36.35 | 2.68 ~23.80 | 2.69 ~ 28.84 | 2.66 ~ 23.86 |
| Map sharpening *B* factor ($Å^2$) | -138.5 | -93.7 | -92.6 | -98.4 |
| Model composition | | | | |
| Non-hydrogen atoms | 9,309 | 9,534 | 9,357 | 9,882 |
| Protein residues | 1,218 | 1,248 | 1,230 | 1,299 |
| Ligands | 15 | 6 | | 6 |
| *B* factors ($Å^2$) | | | | |
| Protein | 60.47 | 47.07 | 52.75 | 51.98 |
| Ligand | 80.93 | 101.74 | | 43.28 |
| R.m.s. deviations | | | | |
| Bond lengths (Å) | 0.736 | 0.968 | 0.534 | 0.533 |
| Bond angles (°) | 0.005 | 0.005 | 0.003 | 0.003 |
| Validation | | | | |
| MolProbity score | 1.36 | 1.22 | 1.25 | 1.17 |
| Clashscore | 4.02 | 4.39 | 4.77 | 3.79 |
| Poor rotamers (%) | 0.00 | 0.00 | 0.00 | 0.00 |
| Ramachandran plot | | | | |
| Favored (%) | 97.00 | 98.21 | 98.27 | 98.37 |
| Allowed (%) | 3.00 | 1.79 | 1.73 | 1.63 |
| Disallowed (%) | 0.00 | 0.00 | 0.00 | 0.00 |

**Table S3: Cryo-EM data refinement and validation of the minor conformation (single protomer)**

| | EAAT3-X 20 mM L-Glu<br>EMD-26986<br>PDB-8CTD | EAAT3-X 300 mM KCl<br>EMD-26998<br>PDB-8CUD | EAAT3-X 150 mM NMDG-Cl<br>EMD-27001<br>PDB-8CUJ | EAAT3-X 300 mM NaCl<br>EMD-27007<br>PDB-8CV3 |
|---|---|---|---|---|
| **Data collection and processing** | | | | |
| Symmetry imposed | C1 | C1 | C1 | C1 |
| Initial particle images (no.) | 1,490,916<br>(C3-expanded) | 1,214,187<br>(C3-expanded) | 630,909<br>(C3-expanded) | 1,559,571<br>(C3-expanded) |
| Final particle images (no.) | 202,573 protomer | 122,507 protomer | 117,532 protomer | 105,451protomer |
| Map resolution (Å) | 3.43 | 2.94 | 3.04 | 3.04 |
| FSC threshold | 0.143 | 0.143 | 0.143 | 0.143 |
| Map resolution range (Å) | 3.01 ~ 32.03 | 2.67 ~ 34.20 | 2.66 ~ 36.59 | 2.69 ~ 41.69 |
| **Refinement** | | | | |
| Initial model used (PDB code) | 6x2z | 6x2z | 6x2z | 6x2z |
| Model resolution (Å) | 3.66 | 3.15 | 3.29 | 3.31 |
| FSC threshold | 0.5 | 0.5 | 0.5 | 0.5 |
| Model resolution range (Å) | 3.66 ~ 19.55 | 3.15 ~ 18.45 | 3.29 ~ 21.07 | 3.31 ~ 19.27 |
| Map sharpening *B* factor (Å$^2$) | -156.8 | -100.7 | -83.5 | -91.8 |
| Model composition | | | | |
| Non-hydrogen atoms | 3,130 | 3,294 | 3,294 | 3,168 |
| Protein residues | 410 | 433 | 433 | 416 |
| Ligands | 5 | | | 2 |
| *B* factors (Å$^2$) | | | | |
| Protein | 62.17 | 56.42 | 75.4 | 41.04 |
| Ligand | 116.27 | | | 36.34 |
| R.m.s. deviations | | | | |
| Bond lengths (Å) | 0.626 | 0.667 | 0.530 | 0.468 |
| Bond angles (°) | 0.003 | 0.005 | 0.002 | 0.002 |
| Validation | | | | |
| MolProbity score | 1.56 | 1.09 | 1.35 | 1.23 |
| Clashscore | 9.77 | 2.95 | 5.90 | 2.76 |
| Poor rotamers (%) | 0.00 | 0.00 | 0.00 | 0.00 |
| Ramachandran plot | | | | |
| Favored (%) | 97.77 | 98.14 | 97.90 | 97.07 |
| Allowed (%) | 2.23 | 1.86 | 2.10 | 2.93 |
| Disallowed (%) | 0.00 | 0.00 | 0.00 | 0.00 |

**Table S4: pKa-s and solvent accessible surface areas of E374**

| States | OFS-Apo$_{KCl}$ | OFS-Apo$_{NMDG}$ | OFS-$Na^+$ | OFS-Glu | iOFS-$K^+$ | iOFS-Apo | iOFS-$Na^+$ | iOFS$^*$-Glu |
|---|---|---|---|---|---|---|---|---|
| pKa[#] | 9.2 | 10.0 | 7.6 | 8.9 | 5.5 | 6.5 | 7.1 | 8.4 |
| ASA (Å$^2$)[$] | 0.2 | 1.6 | 2.1 | 0.0 | 1.3 | 25.3 | 16.2 | 0.0 |

[#] pKa values calculated using PROPKA for chains A

[$] Solvent accessible surface area (ASA) of E374 calculated using PISA for chains A

**Movies S1: Elevator movement of SLC1:** The elevator movements of the transport domain, with HP1 and HP2 colored blue and red respectively, TMs 3, 6, 7, and 8 colored light green, the scaffold domain TMs 1, 2, 4, and 5 colored tan. hEAAT3g protomers in the outward-facing aspartate-bound state (PDB ID 6x2z) and inward-facing Apo state (IFS, PDB ID 6x3f) were used for movie preparation.

**Movies S2: Comparison between aspartate- and glutamate-bound hEAAT3 transport domains**. Structures of OFS-Asp (PDB ID 6x2z) and iOFS$^{*}$-Glu are superimposed on the cytoplasmic half of the transport domain (residues 314-372 and 442-465). TMs 3 and 6 are removed for clarity. HP1, TMs 7 and 8 are colored gray, and HP2, substrate, M367 in NMD motif, and coordinating residues are colored orange and blue, respectively.
